# Mathematical Modeling of Impacts of Patient Differences on Renin-Angiotensin System and Applications to COVID-19 Lung Fibrosis Outcomes

**DOI:** 10.1101/2022.11.06.515367

**Authors:** Mohammad Aminul Islam, Ashlee N. Ford Versypt

## Abstract

Patient-specific premorbidity, age, and sex are significant heterogeneous factors that influence the severe manifestation of lung diseases, including COVID-19 fibrosis. The renin-angiotensin system (RAS) plays a prominent role in regulating the effects of these factors. Recent evidence shows patient-specific alterations of RAS homeostasis concentrations with premorbidity and the expression level of angiotensin-converting enzyme 2 (ACE2) during COVID-19. However, conflicting evidence suggests decreases, increases, or no changes in RAS peptides after SARS-CoV-2 infection. In addition, detailed mechanisms connecting the patient-specific conditions before infection to infection-induced RAS alterations are still unknown. Here, a multiscale computational model was developed to quantify the systemic contribution of heterogeneous factors of RAS during COVID-19. Three submodels were connected—an agent-based model for in-host COVID-19 response in the lung tissue, a RAS dynamics model, and a fibrosis dynamics model to investigate the effects of patient-group-specific factors in the systemic alteration of RAS and collagen deposition in the lung. The model results indicated cell death due to inflammatory response as a major contributor to the reduction of ACE and ACE2. In contrast, there were no significant changes in ACE2 dynamics due to viral-bound internalization of ACE2. The model explained possible mechanisms for conflicting evidence of patient-group-specific changes in RAS peptides in previously published studies. Simulated results were consistent with reported RAS peptide values for SARS-CoV-2-negative and SARS-CoV-2-positive patients. RAS peptides decreased for all virtual patient groups with aging in both sexes. In contrast, large variations in the magnitude of reduction were observed between male and female virtual patients in the older and middle-aged groups. The patient-specific variations in homeostasis RAS peptide concentrations of SARS-CoV-2-negative patients also affected the dynamics of RAS during infection. The model results also showed that feedback between RAS signaling and renin dynamics could restore ANGI homeostasis concentration but failed to restore homeostasis values of RAS peptides downstream of ANGI. In addition, the results showed that ACE2 variations with age and sex significantly altered the concentrations of RAS peptides and led to collagen deposition with slight variations depending on age and sex. This model may find further applications in patient-specific calibrations of tissue models for acute and chronic lung diseases to develop personalized treatments.

## 1. Introduction

The renin-angiotensin system (RAS) has gained significant attention as alterations in RAS peptides and enzymes directly correlate with patient-specific premorbidity, age, and sex differences in response to COVID-19 lung diseases [1–4]. RAS is important in maintaining physiological processes, including blood pressure and vascular permeability [1]. The cascade of RAS’s local and systemic regulatory processes begins with renin from the kidney and angiotensinogen (AGT) from the liver, which enter the circulation through the blood and migrate to the lungs. The renin activity on AGT causes its conversion to angiotensin I (ANGI). ANGI uses the angiotensin-converting enzyme (ACE), including ACE from the cell surface of lung tissue, to convert to angiotensin II (ANGII). ANGII binds with angiotensin-converting enzyme 2 (ACE2) from the cell surface to form the ACE2·ANGII complex, which converts to angiotensin 1–7 (ANG1–7). ANGII also binds with angiotensin type 1 receptor (AT1R) and forms the ANGII·AT1R complex, which regulates the feedback signaling to the systemic renin production rate. In the subsequent reactions, ANGII binds with angiotensin type 2 receptor (AT2R) and forms the ANGII·AT2R complex, ANGII converts to angiotensin IV (ANGIV) by enzymatic reaction, and ANG1–7 binds with MAS receptor to form the MAS·ANG1–7 complex. The expressions of ACE2 [4] and homeostasis values of RAS peptides [5] are patient-dependent.

Kutz et al. [1] observed reductions in all RAS peptides and renin activity without significant differences in ACE and ACE2 activity in the serum of COVID-19 patients compared with matched controls. Reindl-Schwaighofer et al. [6] also observed a decrease in ANGII in COVID-19 patients compared to healthy individuals. Contrarily, the study reported higher ANGII concentrations in severe COVID-19 compared to non-severe COVID-19. Similarly, Liu et al. [2] reported increased ANGII in critically ill SARS-CoV-2 positive patients compared to non-critically ill SARS-CoV-2 positive patients. Another group observed increased plasma ANGII levels in SARS-CoV-2-infected patients compared to healthy individuals [7]. The up-regulation in ACE2 expression has also been observed in COVID-19 patients [8]. Contrary to all, Kintscher et al. [9] did not observe any significant differences in plasma RAS peptides and ACE2 activity between control and COVID-19 patients. To explain the conflicting data on RAS peptide alteration, Kutz et al. [1] highlighted the differences between their study and that of Liu et al. [2], particularly the selection of the baseline patient groups. The baseline patient group in Kutz et al. [1] was SARS-CoV-2-negative, whereas Liu et al. [2] selected SARS-CoV-2-positive patients with normal ANGII range. In addition to these contradictions and limitations of scarce patient data, detailed mechanisms suitable for explaining the conflicting observations in different patient groups still need to be determined. In a three-year study from the start of the COVID-19 pandemic, Prato et al. [10] highlighted the contradictory patient data on RAS without a direct correlation in the imbalance of RAS pathways during SARS-CoV-2 infection. However, the study suggested that age, sex, comorbidities, treatments, and RAS imbalance at the tissue level may influence the COVID-19 severity and outcome.

Patient-specific premorbidity, age, and sex differences can account for response variations between patient groups. Pendergrass et al. [11] observed variations in RAS with sex differences for hypertensive rats. Xudong et al. [12] reported dramatically reduced ACE2 expression with aging and higher ACE2 content in old female rats than in males. In a recent mathematical study, Bastolla [13] showed a direct correlation between the severity of COVID-19 fatality depending on age and sex across three countries and experimental data for the expression of ACE2 from Xudong et al. [12]. Miesbach [14] reported that male patients were more affected by severe manifestations than female patients and depended on the activation of RAS and pathological roles of ANGII. Other studies suggested higher COVID-19 severity in males due to biological differences in RAS, immune systems, and sex hormones [15, 16]. Downregulation of ACE2 inhibits the protective activity of downstream RAS peptides, resulting in inflammation and fibrosis [17, 18]. RAS regulates fibrosis by directly and indirectly activating and inhibiting latent transforming growth factor beta (TGF-β) [19]. ANGII·AT1R complex in RAS activates latent TGF-β directly or via Smad and ERK/p38/MAPK signaling pathways, whereas ANGII·AT2R complex inhibits TGF-β production. Via *in vitro* experiments, ANGII·AT1R was reported to induce procollagen synthesis from human fetal lung fibroblasts directly through mitogenesis and indirectly through TGF-β [20]. ANGII-induced TGF-β production was also observed in vascular smooth muscle cells [21] and glomerular mesangial cells [22]. In our earlier mathematical modeling work [23], we also identified TGF-β as a critical contributor to COVID-19 fibrosis. TGF-β activates and recruits macrophages and fibroblasts, and fibroblasts are responsible for excess collagen deposition and fibrosis [23–25].

Bleomycin-mouse models are widely used to evaluate the effects of RAS peptides and enzymes in lung fibrosis [26]. Li et al. [27] observed increased ANGII and collagen for ACE2 knockout mice in bleomycin-induced lung fibrosis models. Rey-Parra et al. [28] also ran similar experiments for ACE2 knockout male and female mice and observed increased collagen in both males and females with significantly increased collagen in males compared to females [28]; they also highlighted that a higher AT2R/AT1R ratio in female mice could be a possible explanation for the sex differences in collagen deposition. The bleomycin-induced lung fibrosis mouse model of Kuba et al. [29] showed increased ANGII in the presence of SARS-CoV spike proteins. Another study from the same lab reported that ACE2 knockout mice also increased ANGII [30]. The mouse model of Wang et al. [31] showed exogenous ACE2 attenuates bleomycin-induced lung injury by attenuating TGF-β production and collagen deposition.

Multiple computational and mathematical models have been developed to integrate and quantify some aspects of premorbidity, age, and sex differences in RAS and RAS-mediated fibrosis. Leete et al. [32] developed a mathematical model to investigate the impact of sex differences in RAS, and they identified ANGII·AT1R feedback mechanism as a significant modulator of sex differences. Pucci et al. [33] developed a mathematical model to investigate the pathogenic mechanisms of SARS-CoV-2 and RAS and applied their model for *in silico* testing of potential modulators to restore functionality. Sadria and Layton [34] expanded a RAS model with membrane-bound ACE2, shedding of ACE2, and internalized ACE2; connected that to a damage response model; and used the model to investigate the effects of drugs targeting ACE and AT1R. The mathematical modeling framework of Voutouri et al. [35] connected SARS-CoV-2 infection, RAS, innate and adaptive immune cells, and coagulation cascade to investigate the heterogeneity in treatment response and clinical outcome as a function of patient comorbidities, age, and immune response; the model predicted increases in ANGII in severe COVID-19 patients. The extensions of this model investigated the outcomes of immunomodulatory therapies in diverse patient types, including young, diabetic, older, hyperinflated, hypertensive, and obese, [36] and the external and patient-intrinsic factors in disease progression [37]. Barbiero and Lió [38] developed a computational patient model integrating cardiovascular, RAS, and diabetic processes. They analyzed the effects of age, diabetes, and renal impairment during SARS-CoV-2 infection in a functional context. Pacheco-Marin et al. [39] used a discrete Boolean model of RAS, the kallikrein-kinin system, and inflammation for patients affected by COVID-19 to identify the roles of ACE2 in hypertensive and normotensive phenotypes. Our lab has also previously developed mathematical models of RAS for normal and impaired renal functions [40] and glucose dependency of RAS in diabetic kidney disease [41, 42].

Here, we developed a mathematical model that contributes to understanding patient-group-specific premorbidity, age, and sex differences on RAS during SARS-CoV-2 infection. We connected a RAS model to our earlier agent-based model (ABM) framework for in-host tissue response to COVID-19 in the lung [43] and subsequent fibrosis [23] to investigate and quantify the local and systemic effects of the patient-group-specific premorbidity, age, and sex differences on RAS in the progression of fibrosis due to SARS-CoV-2 infection. We hypothesized that variations in the initial number of ACE2 receptors on the surfaces of lung epithelial cells due to age and sex and variations in homeostasis RAS peptide concentrations due to premorbidity result in significant alterations in RAS dynamics during SARS-CoV-2 infection. We tested this hypothesis *in silico* by propagating these conditions through our COVID-19 RAS fibrosis model, defined in the next section.

## 2. Methods

### 2.1. COVID-19 RAS fibrosis model

The COVID-19 RAS fibrosis model (Fig. 1) consists of three submodels: an ABM in-host COVID-19 lung tissue model to quantify tissue scale dynamics of ACE2 receptor trafficking and inflammatory immune response due to SARS-CoV-2 infection (submodel 1 in Fig. 1), a RAS model to account for patient-group-specific local and systemic changes of RAS peptides and enzymes (submodel 2 in Fig. 1), and a fibrosis model to quantify the effects of immune modulation by dysregulated RAS peptides and systemic contributions to lung fibrosis (submodel 3 in Fig. 1). We refer to the COVID-19 RAS fibrosis model as the “overall model.” Each of the submodels is detailed in turn.

**Fig. 1:**
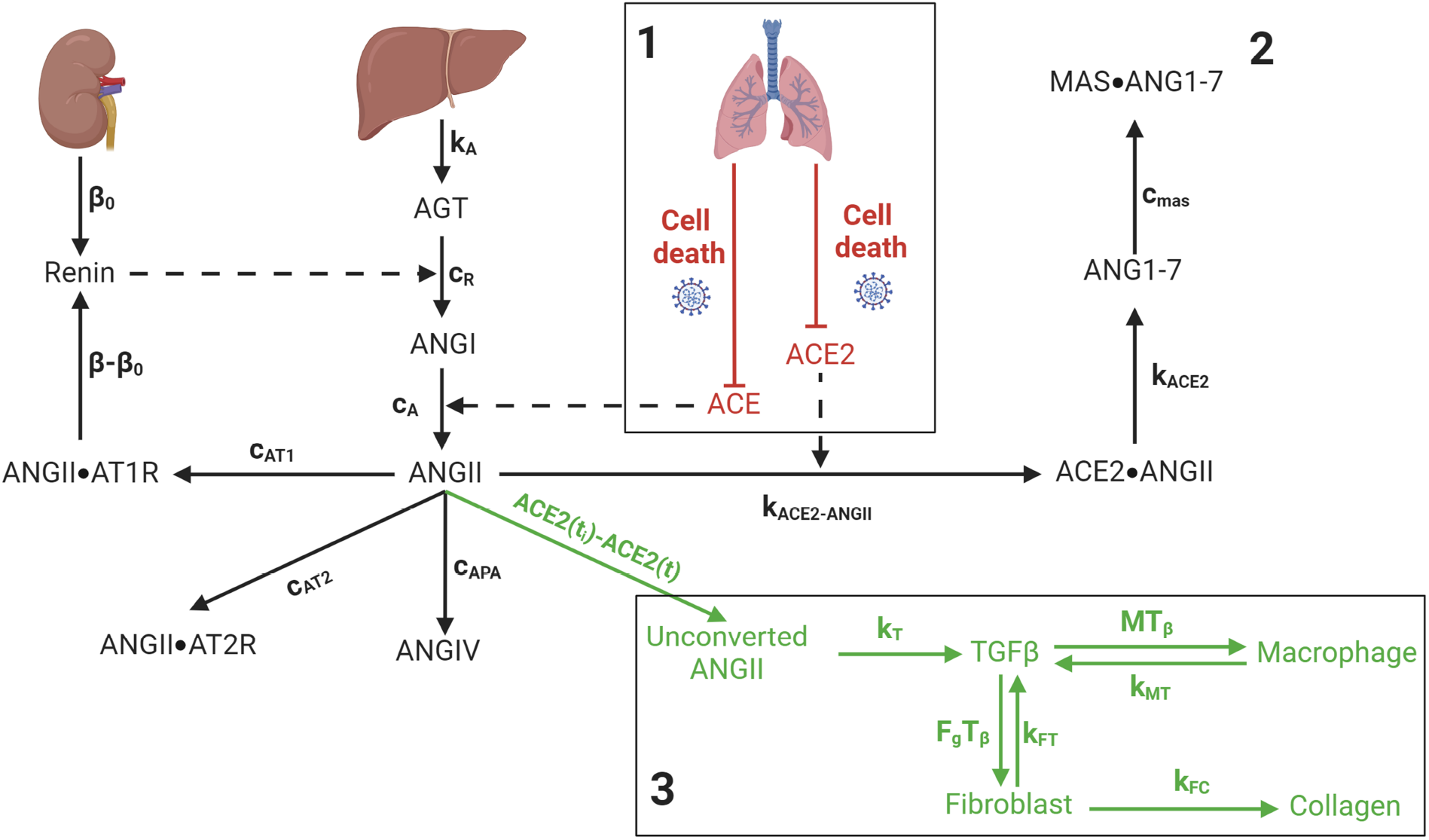
Schematic diagram of COVID-19 renin-angiotensin system (RAS) fibrosis model. Three submodels are denoted by the numbers 1, 2, and 3, respectively, and separated by boxes. The number 1 denotes the agent-based model (ABM) of in-host COVID-19 lung tissue model, number 2 denotes the RAS model, and number 3 denotes the fibrosis model. Renin is produced from the kidney, and angiotensinogen (AGT) is from the liver. The renin activity on AGT causes its conversion to angiotensin I (ANGI). ANGI uses the angiotensin-converting enzyme (ACE) from the cell surface of lung tissue to convert to angiotensin II (ANGII). ANGII binds with angiotensin-converting enzyme 2 (ACE2) from the cell surface to form the ACE2**·**ANGII complex, which converts to angiotensin 1–7 (ANG1–7). ANGII also binds with angiotensin type 1 receptor (AT1R) and forms the ANGII**·**AT1R complex, which regulates the feedback signaling to the systemic renin production rate. In the subsequent reactions, ANGII binds with angiotensin type 2 receptor (AT2R) and forms the ANGII**·**AT2R complex, ANGII converts to angiotensin IV (ANGIV) by enzymatic reaction, and ANG1–7 binds with MAS1 proto-oncogene, G protein-coupled receptor (MAS) to form the MAS**·**ANG1–7 complex. During infection, the death of epithelial cells downregulates both ACE and ACE2, which increases unconverted ANGII. Unconverted ANGII induces transforming growth factor beta (TGF-β) production and activates the fibrosis cascade. Solid arrows denote the transfer of one species to another, and dashed lines denote interactions that influence processes without being produced or consumed. All other notations are defined in Section 2.1 and Tables 2 and 3. Created with BioRender.com.

#### 2.1.1. COVID-19 lung tissue model

In our earlier work, we developed a SARS-CoV-2 tissue simulator [23, 43], which we call the “COVID-19 lung tissue model” (submodel 1 in Fig. 1). The summarized workflow of the agent-based COVID-19 lung tissue model, corresponding equations, and boundary conditions are provided in the Supplementary Material Section S1. Here, we briefly recap the most salient features of the COVID-19 lung tissue model before describing how we use it in this study.

**Table 1:** Age and sex variations in the initial number of unbound external ACE2 receptors per cell (*ueACE*2_0_).

| $ueACE2_0$ | Male | Female |
| --- | --- | --- |
| Older groups |  |  |
| 200 | × |  |
| 400 | × |  |
| 600 |  | × |
| 800 |  | × |
| Middle-aged groups |  |  |
| 1000 | × |  |
| 1200 |  | × |
| Younger groups |  |  |
| 1400 | × |  |
| 1600 | × |  |
| 1800 |  | × |
| 2000 |  | × |

**Table 2:** List of fixed parameters for the overall model.

| Symbol | Definition | Value | Units | Source |
| --- | --- | --- | --- | --- |
| submodel 1 |  |  |  |  |
| $DS$ | TGF-β activation rate from a damaged site | $2 \times 10^{-9}$ | $\text{ng min}^{-1}$ | Estimated in [23] |
| $MS$ | TGF-β secretion rate from an M2 macrophage | $2 \times 10^{-9}$ | $\text{ng min}^{-1}$ | Estimated in [23] |
| submodel 2 |  |  |  |  |
| $c_R$ | Renin activity on AGT | $2 \times 10^1$ | $\text{min}^{-1}$ | [33] |
| $h_R$ | Half-life of renin | $1.2 \times 10^1$ | min | [33, 48] |
| $h_A$ | Half-life of angiotensinogen | $6 \times 10^2$ | min | [33, 48] |
| $h_{A1}$ | Half-life of angiotensin I | $5 \times 10^{-1}$ | min | [33, 48] |
| $h_{A2}$ | Half-life of angiotensin II | $5 \times 10^{-1}$ | min | [33, 48] |
| $h_{A17}$ | Half-life of angiotensin 1-7 | $5 \times 10^{-1}$ | min | [33, 48] |
| $h_{A4}$ | Half-life of angiotensin IV | $5 \times 10^{-1}$ | min | Estimated |
| $h_{AT1}$ | Half-life of angiotensin II & receptor 1 complex | $1.2 \times 10^1$ | min | [33, 48] |
| $h_{AT2}$ | Half-life of angiotensin II & receptor 2 complex | $1.2 \times 10^1$ | min | [33, 48] |
| $h_{mas}$ | Half-life of angiotensin 1-7 & MAS complex | $1.20 \times 10^1$ | min | [33] |
| $\delta$ | Strength of feedback to Renin | $8 \times 10^{-1}$ [0, 1] | Unitless | [32, 33] |
| $[ANGII \cdot AT1R]_L$ | Lower threshold of ANGII-AT1R | 3 | $\text{fmol mL}^{-1}$ | [1] |
| submodel 3 |  |  |  |  |
| $k_T$ | ANGII-induced TGF-β production rate | $5.0 \times 10^{-4}$ | $\text{ng mL min}^{-1} \text{ fmol}^{-1}$ | Estimated |
| $k_{MT}$ | Macrophage TGF-β production rate | $4.86 \times 10^{-5}$ | $\text{min}^{-1}$ | [24, 25] |
| $k_{FT}$ | Fibroblast TGF-β production rate | $2.78 \times 10^{-6}$ | $\text{min}^{-1}$ | [24, 25] |
| $d_T$ | TGF-β degradation rate | $1.04 \times 10^{-2}$ | $\text{min}^{-1}$ | [24] |
| $d_M$ | Macrophage apoptosis rate | $4.17 \times 10^{-4}$ | $\text{min}^{-1}$ | [24, 25] |
| $d_F$ | Fibroblast apoptosis rate | $8.30 \times 10^{-5}$ | $\text{min}^{-1}$ | [24, 25] |
| $k_{FC}$ | Fibroblast collagen production rate | $2.52 \times 10^{-7}$ | $\mu\text{g cell}^{-1} \text{ min}^{-1}$ | Estimated from [49] |
| $V_{T\beta}$ | Rate parameter for TGF-β-dependent collagen deposition | $9.42 \times 10^{-1}$ | Unitless | Estimated in [23] from experimental data of [50, 51] |
| $k_{T\beta}$ | Saturation parameter for TGF-β-dependent collagen deposition | $1.74 \times 10^{-1}$ | Unitless | Estimated in [23] from experimental data of [50, 51] |
| $M(T_\beta)$ | TGF-β-dependent macrophage recruitment rate | interpolated | $\text{cell min}^{-1}$ | [52] |
| $F_g(T_\beta)$ | TGF-β-dependent fibroblast recruitment rate | interpolated | $\text{cell min}^{-1}$ | [52] |

**Table 3:**
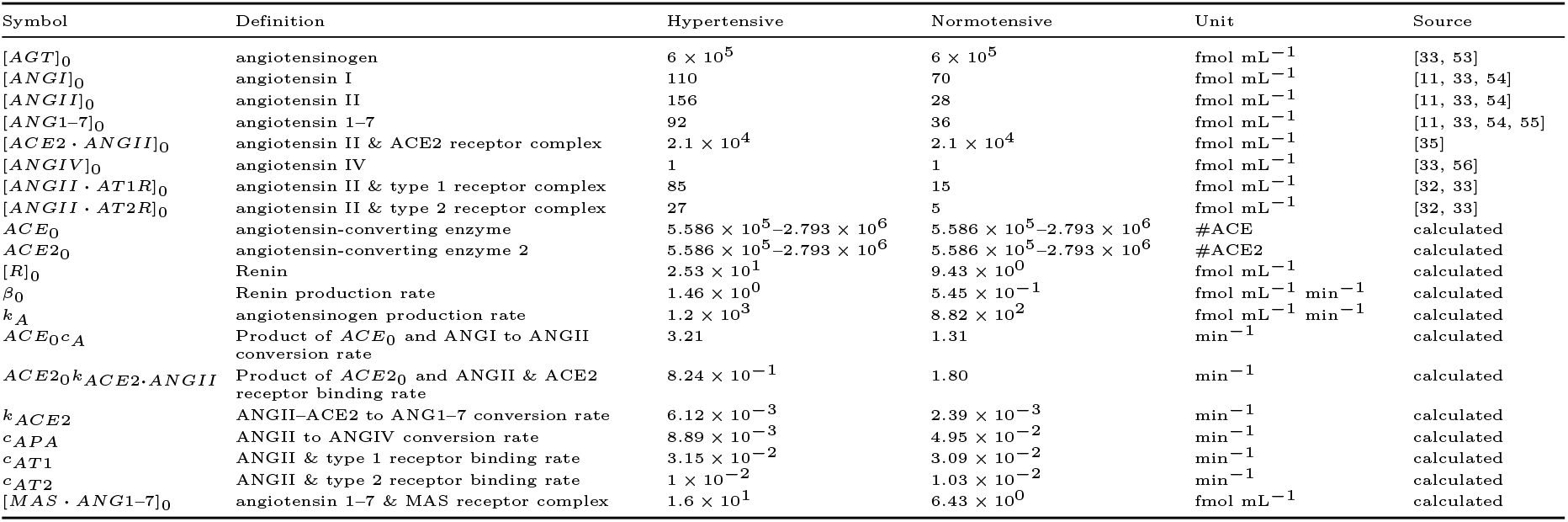
Homeostasis concentrations of species and parameter values in the renin-angiotensin system (RAS) model for uninfected normotensive and hypertensive patients (submodel 2).

The COVID-19 lung tissue model is developed in an open-source multiscale ABM framework PhysiCell [44] and simulates a tissue section of 800 µm *×* 800 µm *×* 20 µm, representing a monolayer of stationary epithelial cells on an alveolar surface of lung tissue. Initially, 2793 epithelial cells, 50 resident macrophages, 28 dendritic cells, and 57 fibroblasts are present in the simulated tissue. Using a uniform random distribution, we infect the simulated tissue by placing SARS-CoV-2 viral particles in the extracellular space. Viral particles diffuse through tissue and bind with unoccupied or unbound external ACE2 receptors (ueACE2) on the epithelial cell surfaces to form bound external ACE2 receptors (beACE2) (Supplementary Material Eqs. (S1)–(S2)). These bound external ACE2 receptors internalize via endocytosis to form bound internal ACE2 receptors (biACE2), release the virions to form unbound internal ACE2 receptors (uiACE2), and recycle back to the cell surface. The released virions from biACE2 replicate through intracellular viral replication kinetics, export back to the extracellular domain by exocytosis, and diffuse in the tissue. Our model considers the interactions between virion and ueACE2 as discrete events, whereas diffusion of the virus in tissue is continuous. Details of the intracellular virus model for replication kinetics, viral response, receptor trafficking, and rules for the discrete to continuous transition of the virus are described in much greater detail in our earlier works [23, 45]. Viral infection activates the innate and adaptive immune response and recruits immune cells from the lymph nodes. Further immune and lymph node model details are available in the Supplementary Material (Eq. (S3)) and elsewhere [23, 43, 45, 46]. Viral infection and immune response activate latent TGF-β and shift the phenotype of pro-inflammatory M1 macrophages to anti-inflammatory M2 macrophages to produce TGF-β in the later phase of infection, which leads to fibroblast-mediated collagen deposition and fibrosis (Supplementary Material Eqs. (S4)–(S10)). Details of the effects of TGF-β sources in fibroblast-mediated collagen deposition at damaged sites of SARS-CoV-2 infected tissue are available in our earlier manuscript [23]. Here, we build upon that work to consider the effects of ACE2 receptor dynamics on the RAS network and lung fibrosis.

In our earlier COVID-19 lung tissue model [23], we considered a single value for the initial number of unbound external ACE2 per epithelial cell (*ueACE*2_0_ = 1000). Here to account for age and sex differences, we vary *ueACE*2_0_ in the range of 200–2000 receptors per cell with a discrete interval of 200. We select the minimum and maximum of the *ueACE*2_0_ values based on those values that generate severe and mild, respectively, infected phenotypes of tissue damage dynamics with the COVID-19 lung tissue model (see results described in Section 3.1). Recent studies showed a direct correlation between ACE2 expression and COVID-19 fatality where aging increased disease severity and male patients were more affected compared to female patients [13, 14]. Thus, we set the *ueACE*2_0_ values for the age and sex of our virtual patient cohort so that lower ranges of *ueACE*2_0_ represent older adults (200–800), intermediate ranges represent middle-aged adults (1000–1200), and upper ranges represent younger adults (1400–2000). The variations of *ueACE*2_0_ within a specific age group are intended to account for sex differences. ACE2 expression in the female rat lung was higher than in male rats [12, 13]. So within a specific age group, we select that lower *ueACE*2_0_ represents males, and higher *ueACE*2_0_ represents females. The age and sex groups with discrete *ueACE*2_0_ are listed in Table 1.

We assume the same initial number of ACE and ACE2 receptors. This assumption is based on the experimental observation of similar intensity of ACE and ACE2 protein expressions in lung tissue in the control mice experiments of Roca-Ho et al. [47]. Losses of ACE and ACE2 can occur due to epithelial cell death and binding between virus and ACE2 during SARS-CoV-2 infection. The simulated results from our tissue model suggested that loss of ACE2 mainly occurs due to cell death after the infection. Compared to that, loss of ACE2 due to viral binding is negligible due to the recycling of ACE2 receptors (details in Section 3.1). Since ACE2 and ACE receptors are present in the cell surface and changes in ACE2 occur due to cell death, we assume that changes in ACE and ACE2 are the same. We run the COVID-19 lung tissue model for variations in the initial values of unbound external ACE2 per epithelial cell (*ueACE*2_0_) to predict age and sex differences in the virtual cohort. The model output of interest from submodel 1 is the tissue-wide dynamic profile for the total number of ueACE2 receptors in the virtual lung tissue. The profile is considered to account for the changes in ACE and ACE2, i.e.,

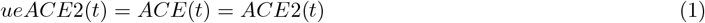

The profiles resulting from the set of initial conditions simulated are passed to the next submodel for the RAS dynamics.

#### 2.1.2. RAS model

We developed a RAS model to account for patient-group-specific local and systemic changes of RAS peptides and enzymes during SARS-CoV-2 infection (submodel 2 in Fig. 1). In the model, the rate of change of renin is

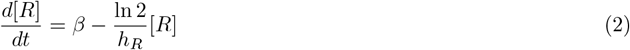

where [*R*] is renin concentration, *β* is the production rate, and *h*_*R*_ is the half-life of renin. The term *β* has two contributions: a constant source of renin from the kidney, *β*_0_, and feedback of ANGII·AT1R to the production of renin, given by

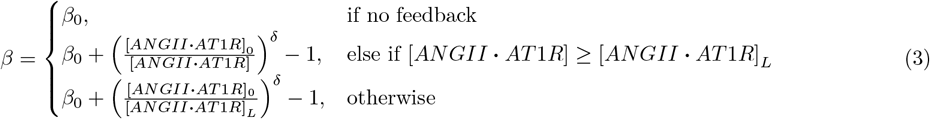

The feedback signaling depends on the initial concentration of ANGII·AT1R ([*ANGII* · *AT* 1*R*]_0_), dynamics of ANGII·AT1R after infection ([*ANGII* · *AT* 1*R*]), and the lower threshold of ANGII·AT1R ([*ANGII* · *AT* 1*R*]_*L*_). We estimated [*ANGII* · *AT* 1*R*]_*L*_ based on the experimental study of Kutz et al. [1]. The strength of the feedback signaling is defined by a dimensionless number, *δ*. The parameter values are listed in Tables 2 and 3.

The rate of change of AGT is

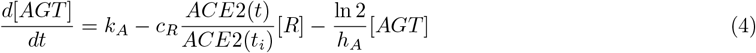

where [*AGT*] is concentration of AGT, *k*_*A*_ is a constant source of AGT from the liver, *c*_*R*_ relates renin concentration to its activity on AGT, *ACE*2(*t*) is the ACE2 receptor tissue-wide dynamic profile determined from the COVID-19 lung tissue model (Section 2.1.1 and Eq. (1)), *ACE*2(*t*_*i*_) is the number of available ACE2 before time of infection (*t < t*_*i*_), and *h*_*A*_ is the half-life of AGT. Here, we consider plasma renin activity (*c*_*R*_) based on earlier models [33, 48]. Kutz et al. [1] reported 58.5% reduction in plasma renin activity in COVID-19 patients. We also assume that changes in the *c*_*R*_ scale are proportional to the ratio of changes in *ACE*2(*t*) in the COVID-19 lung tissue model to account for the changes in plasma renin activity.

The rate of change of ANGI is

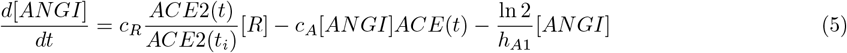

where [*ANGI*] is the concentration of ANGI, *c*_*A*_ is the rate constant for ACE catalyzed conversion of ANGI to ANGII, *ACE*(*t*) is the ACE receptor tissue-wide dynamic profile determined from the COVID-19 lung tissue model (Section 2.1.1 and Eq. (1)), and *h*_*A*1_ is the half-life of ANGI.

The rate of change of ANGII is

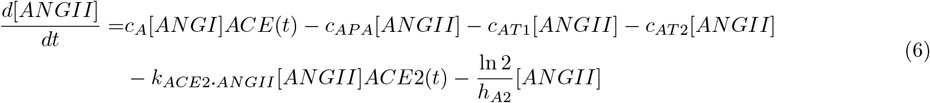

where [*ANGII*] is concentration of ANGII, *c*_*AP A*_ is the rate constant for APA catalyzed conversion of ANGII to ANGIV, *c*_*AT* 1_ and *c*_*AT* 12_ are the binding rate constants for formation of ANGII·AT1R and ANGII·AT2R, respectively, *k*_*ACE*2**·***ANGII*_ is the binding rate constant for formation of ACE2·ANGII, and *h*_*A*2_ is the half-life of ANGII.

The rates of change of ACE2·ANGII and ANG1–7 are

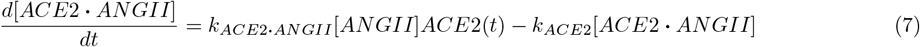

and

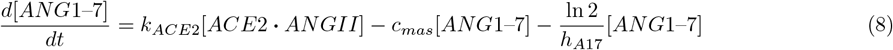

where [*ACE*2 · *ANGII*] and [*ANG*1–7] are concentrations of ACE2·ANGII and ANG1–7, respectively, *k*_*ACE*2_ is the rate constant for the conversion rate of ACE2·ANGII to ANG1–7, *c*_*mas*_ is the binding rate constant for the formation of MAS·ANG1–7, and *h*_*A*17_ is the half-life of ANG1–7. Eqs. (7) and (8) are adapted from the mathematical model of Voutouri et al. [35]. Here, we consider the binding dynamics and reduction in ACE2 from the lung tissue model (Section 2.1.1) and no natural degradation of ACE2·ANGII.

The rate of change of ANGIV is

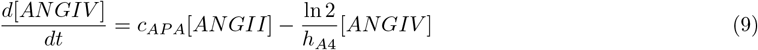

where [*ANGIV*] is concentration of ANGIV and *h*_*A*4_ is the half-life of ANGIV.

ANGII binds with AT1R and AT2R receptors, and ANG1–7 binds with MAS receptors. The rates of change of these complexes are

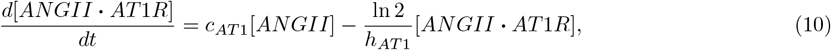

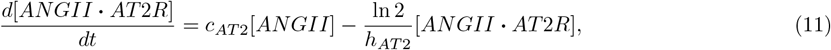

and

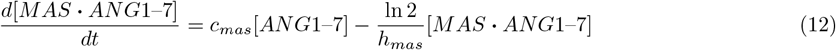

where [*ANGII* · *AT* 1*R*], [*ANGII* · *AT* 2*R*], and [*MAS* · *ANG*1–7] are concentrations and *h*_*AT* 1_, *h*_*AT* 2_, and *h*_*mas*_ are the half lives of ANGII·AT1R, ANGII·AT2R, and MAS·ANG1–7, respectively.

Literature values inform the fixed parameters for submodel 2 (Table 2) and homeostasis RAS peptide concentrations for normotensive and hypertensive patients (Table 3). The parameters [*R*]_0_, *β*_0_, *k*_*A*_, *c*_*A*_, *k*_*ACE*2**·***ANGII*_, *k*_*ACE*2_, *c*_*APA*_, *c*_*AT*1_, *c*_*AT*2_, and [*MAS* · *ANG*1–7]_0_ in Eqs. (2)–(12) are patient-group-specific and require homeostasis values of RAS peptides and enzymes for calibration. We calculate the initial values of ACE (*ACE*_0_) and ACE2 (*ACE*2_0_) by multiplying the number of receptors per cell (*ueACE*2_0_) by the initial number of epithelial cells (2793 cells) from submodel 1. *ACE*(*t*_*i*_) and *ACE*2(*t*_*i*_) are same as *ACE*_0_ and *ACE*2_0_, respectively, before infection. We use the values for *ACE*_0_ and *ACE*2_0_, homeostasis RAS peptide concentrations for normotensive and hypertensive patients (Table 3), and the fixed parameters for submodel 2 (Table 2) to calculate the remaining parameters. To do so, we assume *c*_*mas*_ = *c*_*AT* 2_ and solve Supplementary Material Eqs. (S11)–(S20) for the patient-group-specific parameters (Table 3). The products *ACE*_0_*c*_*A*_ and *ACE*2_0_*k*_*ACE*2**·***ANGII*_ are constant for all *ueACE*2_0_. We use the changing *ACE*_0_ and *ACE*2_0_ values to calculate *c*_*A*_ (#ACE^−1^ min^−1^) and *k*_*ACE*2**·***ANGII*_ (#ACE2^−1^ min^−1^).

Submodel 2 is run for one day to show the homeostasis of RAS peptide concentrations before infection. Then for infection, we use the ACE2 dynamics from submodel 1 as input in submodel 2 to simulate the dynamics of RAS species. Submodel 2 is simulated for ten days post-infection, and ACE2 reaches a new steady state around six days after infection.

#### 2.1.3. Fibrosis model

We developed a fibrosis model to quantify the effects of immune modulation by dysregulated RAS peptides and systemic contributions to lung fibrosis (submodel 3 in Fig. 1). With less ACE2 in the lung, systemic ANGII is converted to ACE2·ANGII at a lower rate, providing some surplus of ANGII not converted along this pathway towards ANG1–7 and increasing the systemic concentration of ANGII. We term the additional ANGII due to the loss of ACE2 as “unconverted ANGII” and calculate the unconverted ANGII as

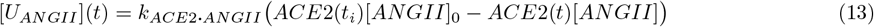

where [*U*_*ANGII*_] is the concentration of unconverted ANGII and the quantities on the right-hand side of Eq. (13) are from submodels 1 and 2 (Sections 2.1.1 and 2.1.2). *k*_*ACE*2**·***ANGII*_ is the binding rate of ANGII to ACE2, [*ANGII*]_0_ is the homeostasis concentration of ANGII, and *ACE*2(*t*_*i*_) is the number of available ACE2 before time of infection (*t < t*_*i*_), which is set to *ACE*2_0_. The values for these three quantities are available in Table 3. *ACE*2(*t*) is the number of available ACE2 receptors during infection (*t* ≥ *t*_*i*_), which is obtained from the tissue-wide dynamic profile from submodel 1 (Section 2.1.1 and Eq. (1)). [*ANGII*] is the concentration of ANGII from submodel 2 (Eq. (6)).

We assume that the unconverted ANGII modulates the direct and indirect activation of TGF-β, and we consider first-order reaction kinetics for ANGII-induced TGF-β production rate. Eq. (14) describes the dynamics of TGF-β:

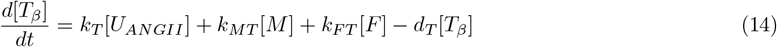

where [*T*_*β*_] is TGF-β concentration, *k*_*T*_ is ANGII-induced TGF-β production rate, *k*_*MT*_ is TGF-β production rate from macrophages, [*M*] is the population of macrophages, *k*_*F T*_ is TGF-β production rate from fibroblasts, [*F*] is the population of fibroblasts, and *d*_*T*_ is degradation rate of TGF-β.

The population balances for macrophages and fibroblasts are

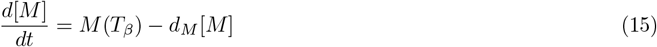

and

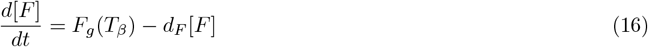

where *d*_*M*_ and *d*_*F*_ are the apoptosis rates of macrophages and fibroblasts, and the TGF-β-dependent macrophage recruitment rate (*M* (*T*_*β*_)) and fibroblast recruitment rate (*F*_*g*_(*T*_*β*_)) are estimated from the experimental observations of Wahl et al. [52]. Linear interpolation is used for the missing ranges of data. Supplementary Material Fig. S1 shows the dynamics of *M* (*T*_*β*_) and *F*_*g*_(*T*_*β*_) with TGF-β.

The TGF-β-dependent collagen deposition rate from fibroblasts is

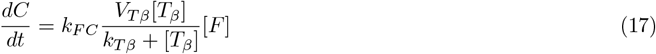

where *C* is the amount of collagen, *k*_*F C*_ is the collagen production rate from fibroblasts, and *V*_*T β*_ and *k*_*T β*_ are corresponding rate and saturation parameters defining the TGF-β dependency on collagen deposition [23].

In the fibrosis model, we use the same parameters as our earlier work [23], except parameter values for TGF-β activation rate from a damaged site (*DS*) and TGF-β secretion rate from an M2 macrophage (*MS*) are fixed (Table 2). Here, the initial number of ACE2 receptors per cell is varied (*ueACE*2_0_, Table 4) to represent virtual patient groups. The initial values of the COVID-19 lung tissue model and fibrosis model variables (Table 4) are all relative to changes from the baseline of an uninfected patient rather than absolute numbers and concentrations.

**Table 4:** Initial conditions for COVID-19 lung tissue and fibrosis submodels.

| Symbol | Definition | Initial condition | Unit |
| --- | --- | --- | --- |
| submodel 1 |  |  |  |
| $ueACE2_0$ | Unoccupied external ACE2 receptors per cell | 200–2000 | # |
| $beACE2_0$ | Virus-bound external ACE2 receptors per cell | 0 | # |
| $biACE2_0$ | Virus-bound internal ACE2 receptors per cell | 0 | # |
| $ueACE2_0$ | Unoccupied internal ACE2 receptors per cell | 0 | # |
| submodel 3 |  |  |  |
| $[T\beta]_0$ | Concentration of TGF-β | 0 | ng mL<sup>-1</sup> |
| $[M]_0$ | Number of macrophages | 0 | cells |
| $[F]_0$ | Number of fibroblasts | 0 | cells |
| $C_0$ | Amount of collagen | 0 | μg |

### 2.2. Sensitivity analysis

We use dynamic local sensitivity analysis for submodel 2 parameters (Section 2.1.2) to quantify the input parameters that significantly affect the output variables. We evaluate the changes in output variables with respect to the one-at-a-time variation of each model parameter. Nominal input value (*I*_*k*_) is the value of the parameter of index *k* calculated using patient-group-specific homeostasis concentrations of RAS peptides (Table 3). The nominal output value (*O*_*k*_(*t*)) is the concentration of a RAS species at time *t* using the parameters in Tables 2 and 3. Each input parameter is multiplied by a multiplier (*m*_*S*_) to change one input at a time. We calculate the change of new input value (*I*_*k,n*_) as Δ*I* = *I*_*k,n*_ − *I*_*k*_, where *I*_*k,n*_ = *m*_*S*_*I*_*k*_. The equation for normalized sensitivity index (*S*_*k,t*_) for the corresponding predicted new output value (*O*_*k,n*_(*t*)) is

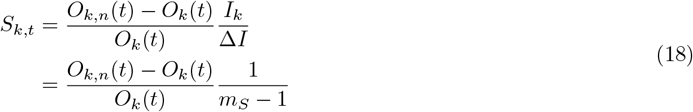

where *k* denotes parameters or variables and *t* denotes time. We apply an additional case to avoid division by zero, as described in Eq. (19):

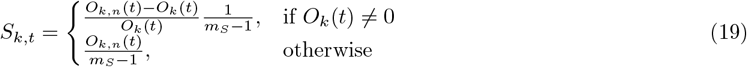

### 2.3 Data extraction

Experimental data from the literature used in this paper were extracted from graphs in the sources (referenced in the text) using the web-based program WebPlotDigitizer [57].

### 2.4. Computational implementation

All the simulations were performed in a Dell Precision 3640 tower workstation: Intel Core Processor i9–10900K (10 core, 20 MB cache, base 3.7 GHz, up to 5.3GHz, and 32GB RAM) using hyperthreading for six total execution threads. For the COVID-19 lung tissue model (submodel 1 in Section 2.1.1), we used PhysiCell (Version 1.9.0) [44] with BioFVM [58] to solve the transport equations (Supplementary Material Section S1). For a single run of submodel 1 with 21,600 minutes (15 days) of simulation, the total wall clock time was around 18 minutes. We used Python 3.8 and the odeint function in the scipy library to solve the differential equations in the RAS model and fibrosis model (submodels 2 and 3 in Sections 2.1.2 and 2.1.3). The code for our earlier fibrosis model is available in a repository at https://github.com/ashleefv/covid19fibrosis [59]. The code for the COVID-19 RAS fibrosis model is available in a repository at https://github.com/ashleefv/covid19fibrosisRAS [60].

## 3. Results and Discussion

The overall model has three submodels, which we analyzed sequentially. First, we infected the virtual lung tissue considering different initial values of unbound external ACE2 receptors per epithelial cell (*ueACE*2_0_) using the COVID-19 lung tissue model (submodel 1 in Section 2.1.1). The COVID-19 lung tissue model was used to evaluate the tissue-wide dynamics of ACE2 receptors after SARS-CoV-2 infection (Section 3.1). Second, we calibrated the RAS model (submodel 2 in Section 2.1.2) with patient-group-specific homeostasis peptide concentrations from Table 3 and an initial number of ACE2 receptors from the COVID-19 lung tissue model (submodel 1). Starting from the homeostasis RAS peptide concentrations before infection, the predicted tissue-wide ueACE2 dynamics during infection from submodel 1 were used as inputs for ACE and ACE2 dynamics (Eq. (1)) in submodel 2. In Section 3.2, we quantified the impacts of age and sex on RAS peptide concentrations predicted by submodel 2 by varying *ueACE*2_0_. Section 3.3 investigated the effects of patient-specific homeostasis concentrations of RAS peptides on submodel 2 results. We quantified the effects of patient-group-specific premorbidity and feedback from downstream RAS signaling to renin by varying initial RAS peptides concentration in Section 3.4. Third, we used the fibrosis model (submodel 3 in Section 2.1.3) to investigate the effects of patient differences in RAS peptides on COVID-19 lung fibrosis outcomes in Section 3.5. Finally, we analyzed the sensitivity of the RAS model to the parameters (Section 3.6) and discussed limitations (Section 3.7).

### 3.1. Dynamics of ACE2 and tissue damage after SARS-CoV-2 infection in response to initial number of ACE2 receptors

In submodel 1, the lower values of *ueACE*2_0_ resulted in higher numbers of infected cells at the earlier phase of infection (*t <* 5 days, Fig. 2A, E, I and Fig. S2). A lower number of *ueACE*2_0_ yields a small binding flux (Eq. (S2)) of virions. As a result, fewer virions bind with an individual cell, and the remaining unbound virions diffuse in the tissue to infect neighboring cells. The virions replicate inside the infected cells, exocytose, diffuse in the tissue, and infect neighboring cells. Also, the removal of infected cells is delayed as the adaptive immune cells (CD8+ T cells) are activated in the later phase of infection (*t >* 4 days, Fig. S3). With lower *ueACE*2_0_, the cascade of events of lower binding flux, diffusion of a higher number of virions, replications of virions, and delayed adaptive immune response cause a higher number of infected cells in the earlier phase of infection. A higher number of viral-bound external ACE2 (beACE2) is observed with lower *ueACE*2_0_ due to virion replications in the infected cells and a higher virion diffusion in the tissue (Fig. 2C, G, K and Fig. S2).

**Fig. 2:**
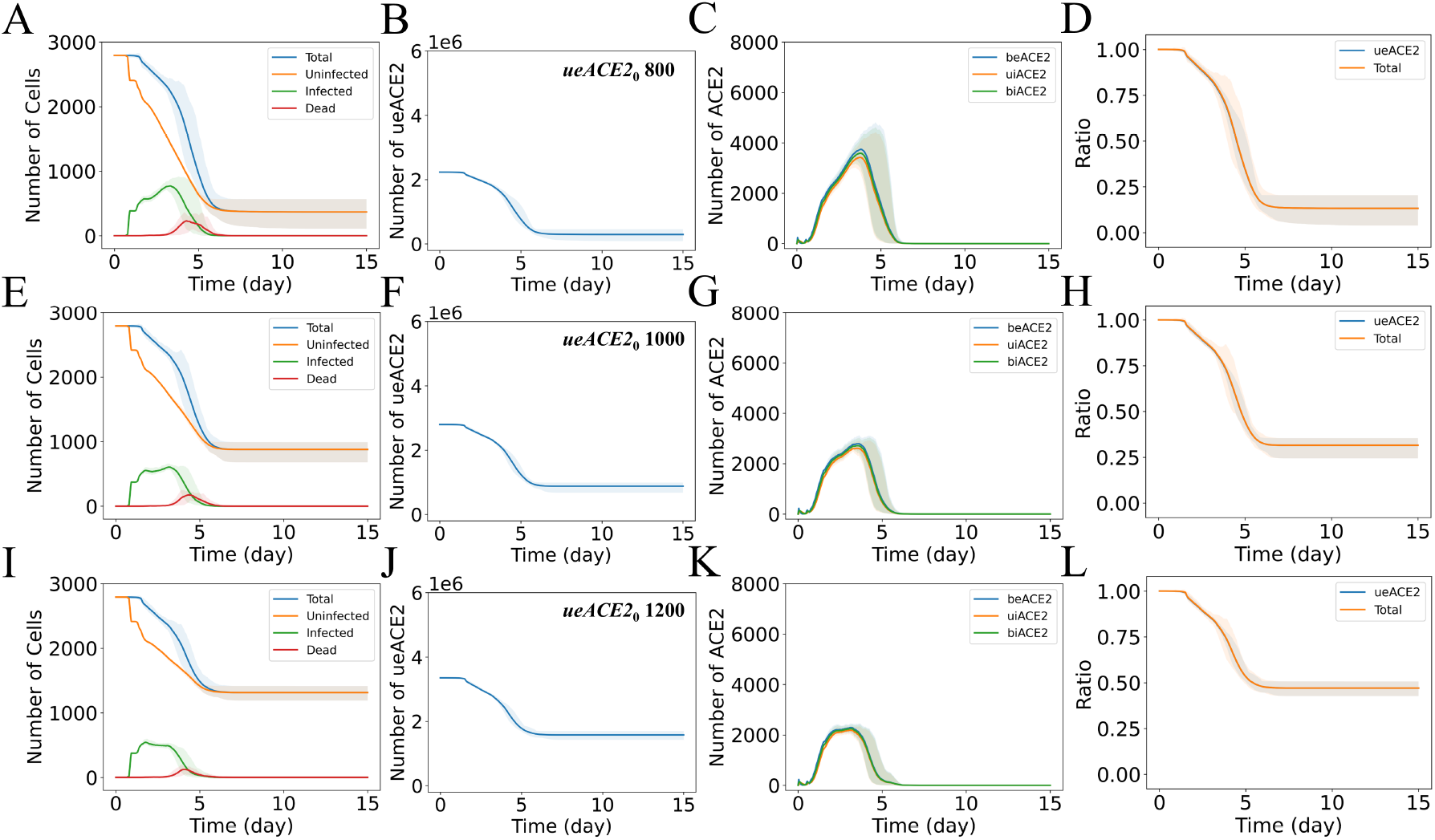
Epithelial cells and ACE2 response to different initial values of unbound external ACE2 per epithelial cell (*ueACE*2_0_). Dynamics of (column 1: A, E, I) total, uninfected, infected, and dead epithelial cells; (column 2: B, F, J) tissue-wide unbound external ACE2 (ueACE2); (column 3: C, G, K) tissue-wide bound external ACE2 (beACE2), bound internal ACE2 (biACE2), and unbound internal ACE2 (uiACE2); and (column 4: D, H, L) normalized comparison between the dynamics of tissue-wide ueACE2 and total cells. Each row represents a fixed *ueACE*2_0_ value in the range of 800–1200 receptors per cell, as labeled in column 2. The numbers of ueACE2 on the *y*-axes of B, F, and J denote the total number of ueACE2 receptors in the virtual lung tissue. The solid curves represent the means, and shaded areas represent the 5th and 95th percentiles of 15 iterations.

We observed variations in the tissue damage dynamics with *ueACE*2_0_. The simulated results for total numbers of epithelial cells (Figs. 2 and S2) showed complete destruction of tissue for *ueACE*2_0_ ≤ 400, increased cell survivability for the range 600 ≤ *ueACE*2_0_ ≤ 1000, and consistent behavior for *ueACE*2_0_ ≥ 1200. The spatial distributions of the epithelial and immune cell populations are shown in Figs. 3 and S3.

**Fig. 3:**
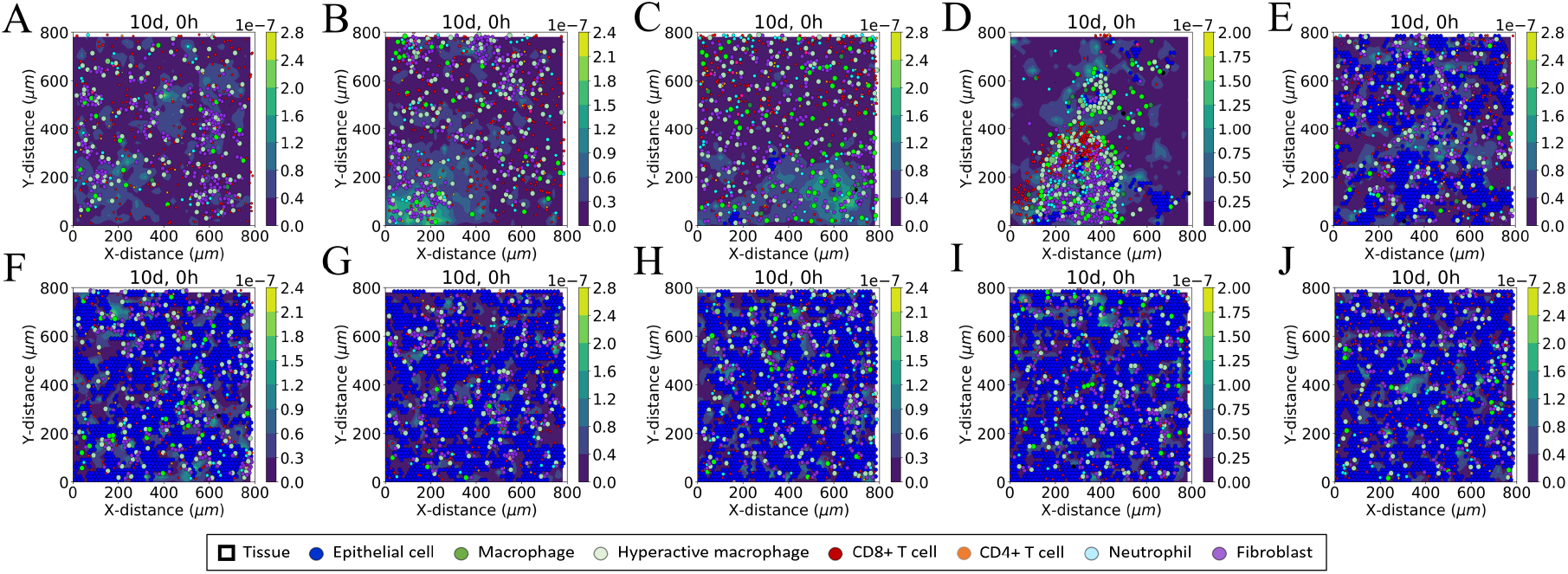
Virtual lung tissue response after 10 days of infection to different initial values of unbound external ACE2 per epithelial cell (*ueACE*2_0_): (A) 200, (B) 400, (C) 600, (D) 800, (E) 1000, (F) 1200, (G) 1400, (H) 1600, (I) 1800, and (J) 2000. Each image is a representative iteration from the set of 15 stochastic iterations of the COVID-19 lung tissue model for each case. Colored circles represent different cell types in the agent-based model (see legend), and the color bars represent the collagen deposited (µg µm^−3^) at damaged sites in tissue.

Figs. 2, S2, and S3 show the dynamics of the epithelial cell populations, tissue-wide ACE2 receptors, and tissue damage with variations in *ueACE*2_0_. The ACE2 receptors reached steady state when the virions were depleted from the system around six days, which corresponds to when the infection dynamics stabilize (Fig. 2A, E, I). While the values of *ueACE*2_0_ are low, there are few available ACE2 receptors, and the steady-state value of ACE2 receptors after infection approaches zero; under these conditions, changing between *ueACE*2_0_ increments substantially impacts the outcomes (Fig. 2B, F, J and Fig. S2).

There were not significant variations in ACE2 dynamics due to viral binding and internalization of ACE2 (beACE2, biACE2, and uiACE2) as the recycling process moved a portion of internalized ACE2 back to the surface. There was ample availability of surface ACE2 receptors in neighboring uninfected cells (Fig. 2C, G, K). However, a direct correlation between the dynamics of total number of epithelial cells and ueACE2 was observed, and the two curves completely overlap when normalized (Fig. 2D, H, L). The findings suggest that the dysregulation of ACE2 in the lung tissue mainly occurs due to the loss of major sources of ACE2 by the death of epithelial cells during infection.

### 3.2. Dynamics of RAS peptides with variations in ACE2 for age and sex differences

We investigated the influence of age and sex in the dynamics of RAS peptides predicted by submodel 2 by varying *ueACE*2_0_, according to the values in Table 1. Here, we considered the submodel 2 outputs of peptide concentrations profiles for the case of Group 1 of our virtual patient groups (defined below in Section 3.4): hypertensive patients with no feedback of ANGII·AT1R to the production of renin. Later, we repeated the analysis for the other virtual patient groups, and the effects are discussed in Section 3.4. Figs. 4 and S4 show the dynamics of RAS peptides. We observed a decrease in ANGI and downstream RAS peptides due to the loss of ACE and ACE2 via inflammatory cell death. We also observed decreased ANGI and downstream peptides for all patient groups with aging in both sex groups. The magnitude of the reduction is largest for older patients (solid curves), intermediate for middle-aged patients (dashed curves), and smallest for younger patients (dotted curves). The percentages of changes in the RAS peptides from the homeostasis concentrations are reported in Supplementary Material Table S1. We observed reductions of 66–100% of ANGI, 93–100% of ANGII, and 99–100% of ANG1–7 for older patients, 25–39% of ANGI, 56–74% of ANGII, and 79–92% of ANG1–7 for middle-aged patients, and 16–19% of ANGI, 40–46% of ANGII, and 63–70% of ANG1–7 for younger patients at day 10 (Fig. 4 and Table S1). These results suggest significant variations in RAS peptides with aging. Clinical studies also reported non-mild COVID-19 cases with aging and higher COVID-19 severity for middle-aged and older patients [61, 62]. So, our findings are consistent with the idea that variations in RAS peptides due to aging can be a factor in a higher susceptibility of severe COVID-19 cases for older and middle-aged patients than for younger patients.

**Fig. 4:**
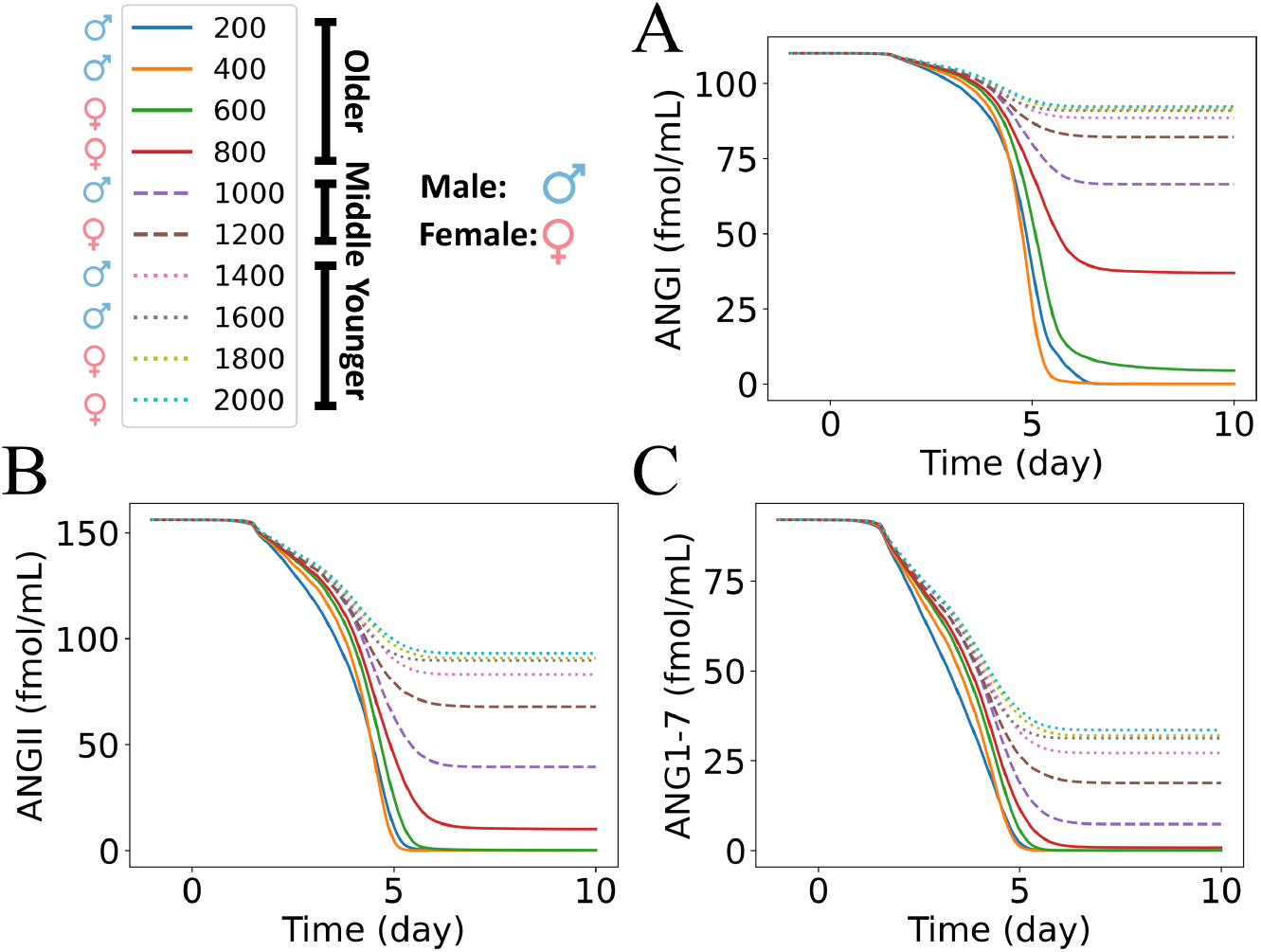
Dynamics of RAS peptides in response to different initial values of unbound external ACE2 per epithelial cell (*ueACE*2_0_) for Group 1: hypertensive patients with no feedback from ANGII**·**AT1R to renin. Dynamics of (A) ANGI, (B) ANGII, and (C) ANG1–7 for *ueACE*2_0_ values in the range of 200–2000 recepters per cell. The legend shows the age and sex labels for each discrete *ueACE*2_0_ value (see also Table 1).

We used discrete values of *ueACE*2_0_ within each age group range to differentiate between males and females (Table 1). The simulated results showed large variations in the magnitude of reduction between male and female patients in the older and middle-aged groups (Table S1). In the older group, males had 100% reduction of ANGI, ANGII, and ANG1–7 due to the complete destruction of the virtual tissue, whereas females had 66–95% of ANGI, 93–99% of ANGII, and 99–100% of ANG1–7 reduction at day 10 (Table S1). The middle-aged male results showed a reduction of 39% of ANGI, 74% of ANGII, and 92% of ANG1–7, whereas the middle-aged female results showed a reduction of 25% of ANGI, 56% of ANGII, and 79% of ANG1–7 (Table S1). In contrast, the variations due to sex were relatively slight for the young patient group. The studies on sex differences in COVID-19 showed male patients had higher susceptibility, severity, and longer length of hospital stay compared to female patients [63, 64]. Our simulated dynamics successfully predicted variations in RAS due to sex, which can be a factor associated with sex disparities in COVID-19 outcomes.

### 3.3. Dynamics of RAS peptides with variations in patient-specific initial values of RAS peptides

The patient data in Kutz et al. [1] showed decreases in ANGI, ANGII, and ANG1–7 for COVID-19 patients. Kutz et al. [1] investigated RAS peptide alteration between SARS-CoV-2-positive and SARS-CoV-2-negative patients with respiratory infections. Their study included older adults with premorbid conditions, such as hypertension, diabetes, and obesity, for both SARS-CoV-2-positive and SARS-CoV-2-negative patients. They observed lower equilibrium serum RAS peptides and plasma renin activity in infected patients compared to non-infected patients. They also reported the ranges of variations in ANGI (3–595 pmol L^−1^), ANGII (10–1687 pmol L^−1^), and ANG1–7 (3–166 pmol L^−1^) in SARS-CoV-2-negative patients and the ranges of variations in ANGI (2.7–188 pmol L^−1^), ANGII (2.1–357 pmol L^−1^), and ANG1–7 (3–7.3 pmol L^−1^) in SARS-CoV-2-positive patients. However, the time points for the data were not reported.

When the initial values of RAS peptides were set to those listed in Table 3, our simulated dynamics of RAS in Fig. 4 also showed a decrease in ANGI, ANGII, and ANG1–7 for all *ueACE*2_0_ values. Here, the effects of variations in the initial values of RAS peptides were investigated. Instead of varying *ueACE*2_0_ values as the input to submodel 1 as in Section 3.2, we considered patient-specific homeostasis values of RAS peptides before infection and compared them to RAS peptide concentrations after 10 days of infection (sampled from the data ranges in Kutz et al. [1]) to evaluate if submodels 1 and 2 could capture the experimentally observed decreases in RAS peptides during SARS-CoV-2 infection. To do so, we sampled 1000 sets of initial values of ANGI, ANGII, and ANG1–7 using log uniform distributions within the ranges of experimentally observed values of SARS-CoV-2-negative patients [1]. We used ANGI ranges of 2–600 fmol mL^−1^, ANGII ranges of 2–1700 fmol mL^−1^, and ANG1–7 ranges of 2–150 fmol mL^−1^.

We accounted for the relative differences in the initial values of ANGI, ANGII, and ANG1–7 by sampling within the bounds:

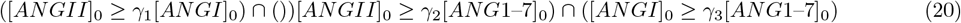

where *γ*_1_ is the ratio of ANGII to ANGI, *γ*_2_ is the ratio of ANGII to ANG1–7, and *γ*_3_ is the ratio of ANGI to ANG1–7. We selected *γ*_1_ = 1.4, *γ*_2_ = 1.7, and *γ*_3_ = 1.2 using the homeostasis concentrations of RAS peptides for hypertensive patients listed in Table 3. ANGIV, ANGII·AT1R, and ANGII·AT2R peptides are derived from ANGII. So, the variations in the initial values of ANGII also affect the initial values of ANGIV, ANGII·AT1R, and ANGII·AT2R. The concentrations of ANGIV, ANGII·AT1R, and ANGII·AT2R were scaled based on ANGII value for hypertensive patients listed in Table 3, and new ANGII values were sampled from the distribution. For a sampled ANGII value ([*ANGII*]_0,*s*_) and ANGII value for hypertensive patients listed in Table 3 ([*ANGII*]_0_), we calculated the scaling factor *S*_*f*_ as

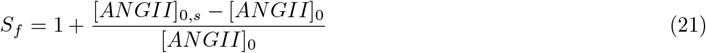

Then, the homeostasis concentrations of [*ANGIV*]_0_, [*ANGII* · *AT* 1*R*]_0_, and [*ANGII* · *AT* 2*R*]_0_ for hypertensive patients from Table 3 were multiplied with *S*_*f*_ (Eq. (21)). We used the dynamics of ACE and ACE2 from submodel 1 for *ueACE*2_0_ = 1000 and followed the methods in Section 2.1.2 as when the homeostasis RAS peptide concentrations from Table 3 were used. Here, the sampled RAS peptide concentrations and calculated *ACE*_0_ and *ACE*2_0_ from *ueACE*2_0_ = 1000 were used to solve Supplementary Material Eqs. (S11)–(S20) and update the parameters ([*R*]_0_, *β*_0_, *k*_*A*_, *c*_*A*_, *k*_*ACE*2**·***ANGII*_, *k*_*ACE*2_, *c*_*APA*_, *c*_*AT*1_, *c*_*AT*2_, and [*MAS* · *ANG*1–7]_0_) for each virtual patient sample.

With the updated parameters and virtual patient homeostasis concentrations of RAS peptides before infection, we used submodel 2 to predict the dynamics of RAS peptides. We compared the simulated ranges of ANGI, ANGII, and ANG1–7 from 1000 virtual patients to the ranges of SARS-CoV-2-positive patients [1] (Fig. 5). Fig. 5A–C show the resulting dynamics of ANGI, ANGII, and ANG1–7. Fig. 5D–F show the subsets of the curves from Fig. 5A–C where the reductions in ANGI, ANGII, and ANG1–7 were all in similar ranges as the experimentally observed values of SARS-CoV-2-positive patients at ten days post-infection. Our simulated results showed the dynamics of 65 virtual patients in Fig. 5D–F that were consistent with experimentally reported ANGI, ANGII, and ANG1–7 values from SARS-CoV-2-negative and SARS-CoV-2-positive patients. The analysis suggests that the patient-specific homeostasis concentrations of RAS peptides before infection affect the dynamics of RAS during infection. Premorbidty, i.e., hypertension, diabetes, chronic kidney disease, and cardiovascular diseases, can alter the homeostasis concentrations of RAS [10]. Thus, it is necessary to include the effects of premorbidity with age and sex to predict disease severity and outcomes of COVID-19 accurately.

**Fig. 5:**
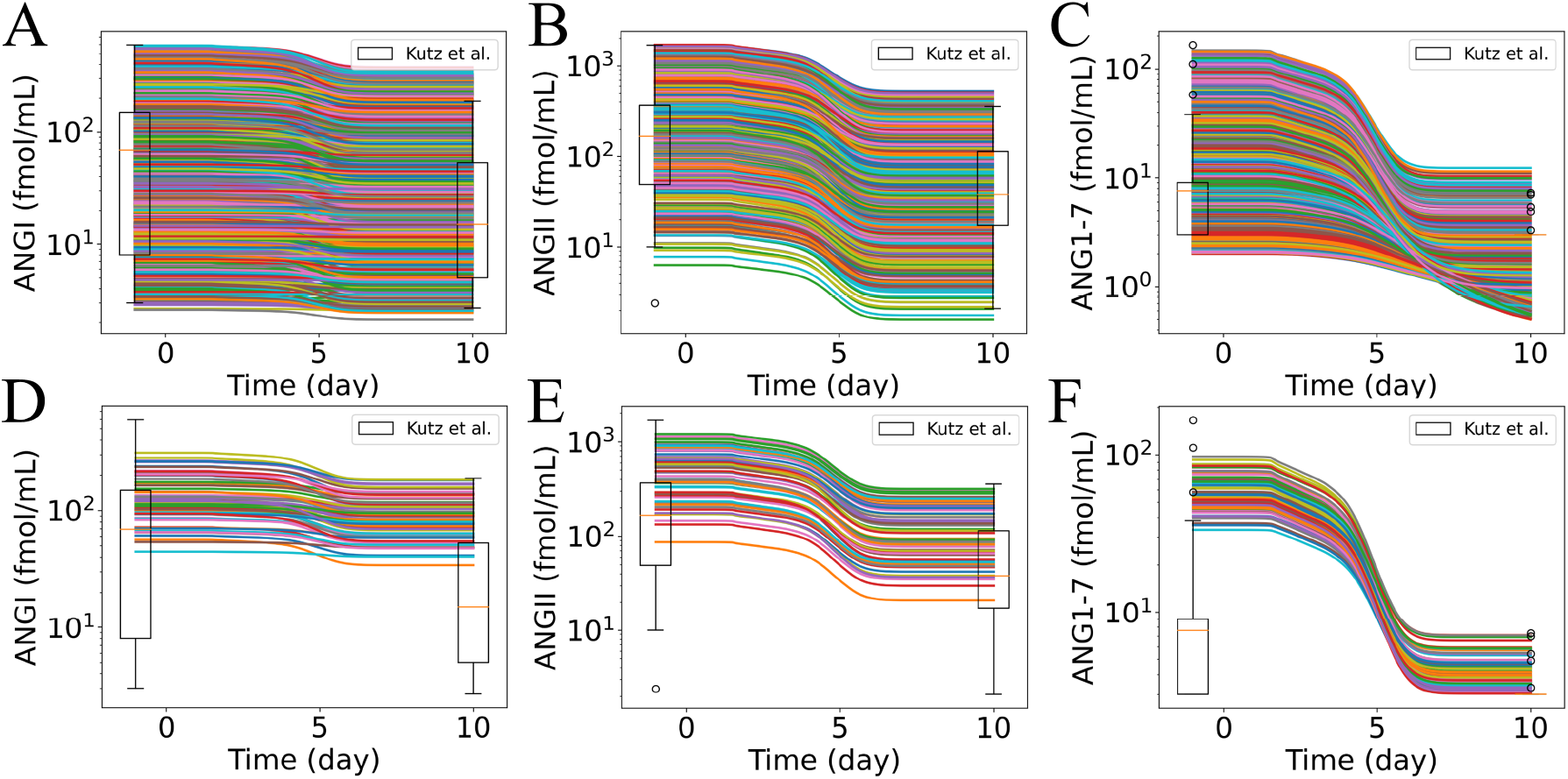
Effects of initial sampled values of ANGI, ANGII, and ANG1–7 on the dynamics of RAS peptides. (A, B, C) RAS peptide dynamics resulting from the initial values of ANGI, ANGII, and ANG1–7 sampled within the experimentally observed ranges of SARS-CoV-2-negative patients reported in Kutz et al. [1] (shown at one day before infection). (D, E, F) The subset of the curves from the corresponding figure in the top row that show similar reduction within the experimentally observed ranges of values from SARS-CoV-2-positive patients reported in Kutz et al. [1] at ten days post-infection. The subset was selected if simulated dynamics of all three RAS peptides of interest (ANGI, ANGII, and ANG1–7) were within the experimental ranges simultaneously at ten days post-infection.

### 3.4. Effects of premorbidty and feedback from ANGII·AT1R to renin

We investigated the effects of premorbidity by varying homeostasis RAS peptide concentrations (hypertensive and normotensive patients from Table 3) and by molecular dysregulation in the feedback signaling to renin. The feedback signaling from ANGII·AT1R to renin has been shown to influence the dynamics of RAS peptides [32]. However, Neubauer et al. [65] reported no evidence for direct feedback of ANGII·AT1R to renin in the mouse kidney. Instead, they suggested that feedback of ANGII·AT1R to renin originates from *in vitro* experiments; in their *in vivo* settings, the required concentrations of ANGII to induce renin secretion from renin cells were not achieved. As an alternative, the experimental study proposed an indirect effect of ANG II on renin-secreting cells (e.g., blood pressure) [65]. Here, we investigated the effect of feedback signaling as a premorbid condition. We assumed that the absence of feedback signaling is a form of molecular dysregulation. As variations in the strength of the feedback signaling (δ) have previously been used to account for sex effects by Leete et al. [32], we selected a constant value of δ (Table 2) so that in our model, the effects of sex are due to the variations in *ueACE*2_0_. We considered four virtual patient groups for premorbidity and feedback:

- Group 1: hypertensive patients with no feedback from ANGII·AT1R to renin
- Group 2: normotensive patients with no feedback from ANGII·AT1R to renin
- Group 3: hypertensive patients with feedback from ANGII·AT1R to renin
- Group 4: normotensive patients with feedback from ANGII·AT1R to renin

Figs. 4 and S4 show the dynamics of RAS peptides for Group 1, and Fig. S5 shows the dynamic results for all four virtual patient groups for combinations of premorbidity and feedback. As for Group 1 discussed in Section 3.2, Group 2 had decreases in ANGI and downstream peptides for all *ueACE*2_0_ values with time during infection and with aging in both sexes. Groups 3 and 4 had small ANGI changes of only ±2% for *ueACE*2_0_ *>* 800, and the other downstream peptides decreased in a similar manner as for the other groups. The starting magnitudes of RAS peptides for Groups 1 and 3 (hypertensive) were higher, which made the absolute magnitude of their reductions larger than for Groups 2 and 4 (normotensive).

Figs. 6 and S6 show the dose-response of how input values of *ueACE*2_0_ affect submodel 2 responses of percent change in RAS peptides at 10 days post-infection. The percentages of changes in the RAS peptides from the homeostasis concentrations at day 10 for all groups are reported in Tables S1–S4. For the lowest *ueACE*2_0_ values, the percentages of changes are ≈100% reduction in ANGI and downstream peptides. The percentages of changes have similar ranges across groups for most peptides, with the exception of those for renin and AGT. Renin had no changes for Groups 1 and 2 without feedback, and large but different ranges of increases for Groups 3 and 4 with feedback. AGT changes differed between premorbity ranges with Groups 1 and 3 (hypertensive) having dose-responses in the range of 19–73%, and dose-responses in the lower ranges of 4–27% for Groups 2 and 4 (normotensive). For the *ueACE*2_0_ *>* 600 values that were not at the maximum reduction, we observed larger absolute values of changes in ANGI for normotensive patients (Groups 2 and 4) compared to hypertensive patients (Groups 1 and 3) with and without feedback. For the other downstream peptides and *ueACE*2_0_ *>* 800, we observed slightly stronger dose-responses (change in peptide from baseline) in normotensive patients compared to hypertensive patients without feedback (matching colored points shift slightly lower from Group 1 to Group 2 in Figs. 6 and S6, and the values in Table S1 have smaller absolute values than those in Table S2). The opposite effect was observed with feedback; there were substantially weaker dose-responses in normotensive patients compared to hypertensive patients (matching colored points shift higher from Group 3 to Group 4 in Figs. 6 and S6, and the values in Table S3 have larger absolute values than those in Table S4). The results suggest the importance of feedback signaling from ANGII·AT1R to renin. The lower homeostasis RAS peptide concentrations for normotensive patients without feedback signaling in Group 2 contributed to a larger magnitude of reduction. In contrast, lower homeostasis RAS peptide concentrations with feedback signaling in Group 4 made the magnitude of reduction smaller (Tables S2 and S4).

**Fig. 6:**
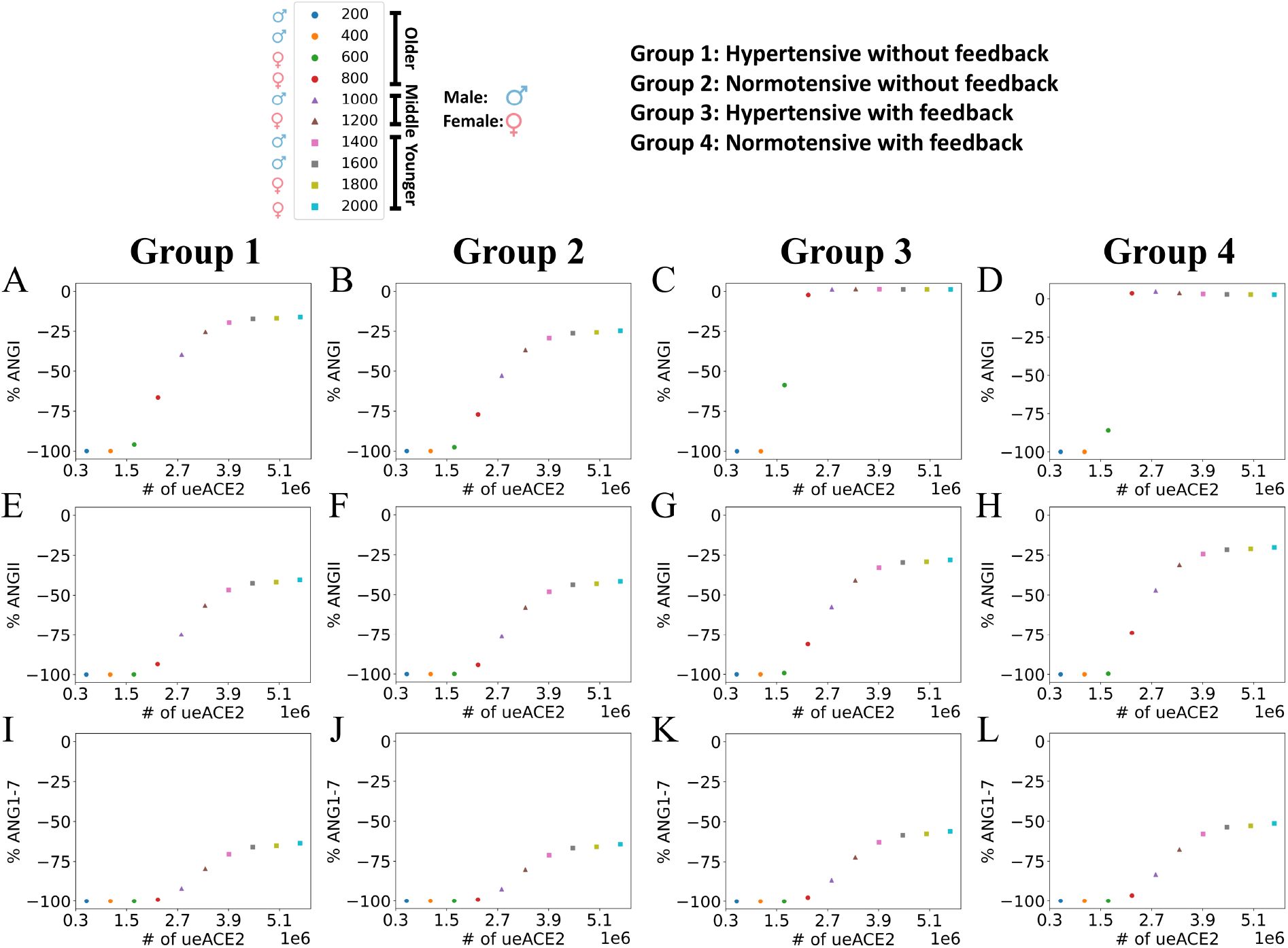
Dose-response at 10 days after infection to different initial values of unbound external ACE2 per epithelial cell (*ueACE*2_0_) for percent change in RAS peptides compared to baseline. (Row 1: A–D) ANGI, (row 2: E–H) ANGII, and (row 3: I–L) ANG1–7. Patient groups represent each column. Group 1 (column 1): hypertensive patients with no feedback from ANGII**·**AT1R to renin. Group 2 (column 2): normotensive patients with no feedback from ANGII**·**AT1R to renin. Group 3 (column 3): hypertensive patients with feedback from ANGII**·**AT1R to renin. Group 4 (column 4): normotensive patients with feedback from ANGII**·**AT1R to renin. The legend shows the age and sex labels for each discrete *ueACE*2_0_ value (see also Table 1).

The feedback of ANGII·AT1R to renin was able to restore the ANGI concentration to the homeostasis level for both hypertensive and normotensive middle-aged and younger patients (Groups 3 and 4 in Fig. 6 and Tables S3 and S4). The feedback signaling to renin (Groups 3 and 4) increased the renin concentrations dynamically compared to the uniform profiles that were independent of *ueACE*2_0_ values for Groups 1 and 2 without feedback. Although the older groups had the largest magnitude of increase in renin concentration, these changes failed to modulate RAS peptide concentration due to the survivability of no or few cells in the tissue at day 10 (*ueACE*2_0_ ≤ 600, Fig. 3A, B, C). However, in the older female (*ueACE*2_0_ = 800), middle-aged, and younger groups, the increase of renin concentrations restored ANGI at day 10 (Group 3 and Group 4 in Fig. S6 and Tables S3 and S4). So, depending on age and sex, feedback of ANGII·AT1R to renin could restore ANGI homeostasis. The molecular dysregulation case of no feedback shows the impact of one change that can drastically alter the ANGI responses in some patient groups. Our simulation results showed that the magnitude of variations of RAS peptides depends on the premorbidity associated homeostasis concentrations of RAS peptides and feedback of ANGII·AT1R to renin. Reindl-Schwaighofer et al. [6] reported a larger magnitude of reduction of ANGII in non-severe COVID-19 patients than in severe COVID-19 patients compared to healthy individuals. However, the study reported a decreased ANGI in non-severe COVID-19 and increased ANGI in severe COVID-19 compared to healthy individuals. Our model is able to reconcile these seemingly contradictory and nonlinear results for some combinations of parameters (see values in Tables S1–S4). For example, consider *ueACE*2_0_ = 800 and Group 4 (older female normotensive with feedback) as the severe case and *ueACE*2_0_ = 1000 and Groups 1 and 2 (middle-aged male hypertensive and normotensive without feedback) as non-severe cases. ANGI for these non-severe cases decreased (%ANGI = -39.65 and -52.81 for Groups 1 and 2, respectively), and ANGI increased for the severe case (%ANGI = 3.58). For the same sets of cases, the non-severe cases reduced ANGII by 74.68% and 76.19% (Groups 1 and 2) compared to a reduction of only 73.84% in the severe case. These results are consistent with the Reindl-Schwaighofer et al. [6] observations. Kutz et al. [1] also observed variations in the reduction of ANGI and ANGII in SARS-CoV-2-positive patients, which are evident in our analysis in Section 3.3.

### 3.5. Dynamics of fibrosis model with patient differences

Multiple studies identified a reduction in ACE2 and an increase in ANGII as regulators of excess collagen deposition and fibrosis [27–31]. Here, we assumed that the reduction of ACE2 in the lung tissue contributes to a surplus of ANGII (unconverted ANGII) at a systemic level and increases the systemic concentration of ANGII. This is the premise for submodel 3. In a study, Liu et al. [2] collected plasma samples and demographic data from 55 SARS-CoV-2-positive patients. They reported increased plasma ANGII above the normal range of plasma ANGII (25–129 pg mL^−1^), mainly in critically ill patients. The measured range of ANGII in the critically ill patients group was 149.7 (137.8, 165.1) pg mL^−1^, where the values represent mean (lower bound, upper bound). So, from the mean of the normal range of ANGII (77 pg mL^−1^), ANGII in the critically ill patients group increased by 94.4% (79%, 114.4%). Here, we assumed that the increase in ANGII in critically ill patients compared to the mean value of ANGII in normal patients is due to the dysregulation of ACE2. Unconverted ANGII in our model represents the difference in ANGII between baseline ACE2 (*ACE*2(*t*_*i*_)) and reduced ACE2 dynamics considering cell death (*ACE*2(*t > t*_*i*_), Eq. (13) in Section 2.1.3).

Fig. 7 shows the dynamic profiles of the percentage of unconverted ANGII increase from baseline with initial RAS peptide concentration variations for the 1000 virtual patients sampled in Section 3.3 at *ueACE*2_0_ = 1000. We predicted a similar increase in unconverted ANGII as the experimental observations of Liu et al. [2] with a mean of 92.12, median of 92.06, and standard deviation of 1.21. The similarities in the profiles (Fig. 7) demonstrate that unconverted ANGII depends more on the dynamics of ACE2 (which are the same for all from submodel 1 with *ueACE*2_0_ = 1000) than on the initial values of the RAS peptides that vary for the 1000 virtual patients. For the four virtual patient groups defined in Section 3.4 and varying *ueACE*2_0_, Fig. 8A–D show the dynamics of unconverted ANGII, and Table S5 tabulates the percentage increase in unconverted ANGII at day 10 post-infection. In our model, the differences between hypertensive (Group 1 and Group 2) and normotensive (Group 2 and Group 4) groups are the homeostasis RAS peptide concentrations. The largest effects were due to variations in *ueACE*2_0_ (curves on the same plot in Fig. 8A–D compared to panels for each patient group). Similar to Fig. 7, we observed slight variations in unconverted ANGII between hypertensive and normotensive groups without feedback (Group 1 and Group 2) and with feedback (Group 3 and Group 4) for a specific *ueACE*2_0_ value (Table S5). However, the presence of feedback signaling also affected the dynamics of unconverted ANGII. In the absence of feedback signaling, Group 1 showed an increase in unconverted ANGII of 99–100% for older patients, 79–92% for middle-aged patients, and 63–70% for younger patients. For Group 3, the increases in unconverted ANGII were 97–100% for older, 72–87% for middle-aged, and 56–63% for younger patients (Table S5). From the observations, we predicted that age, sex, and feedback signaling from ANGII·AT1R to renin are important factors in regulating systemic ANGII.

**Fig. 7:**
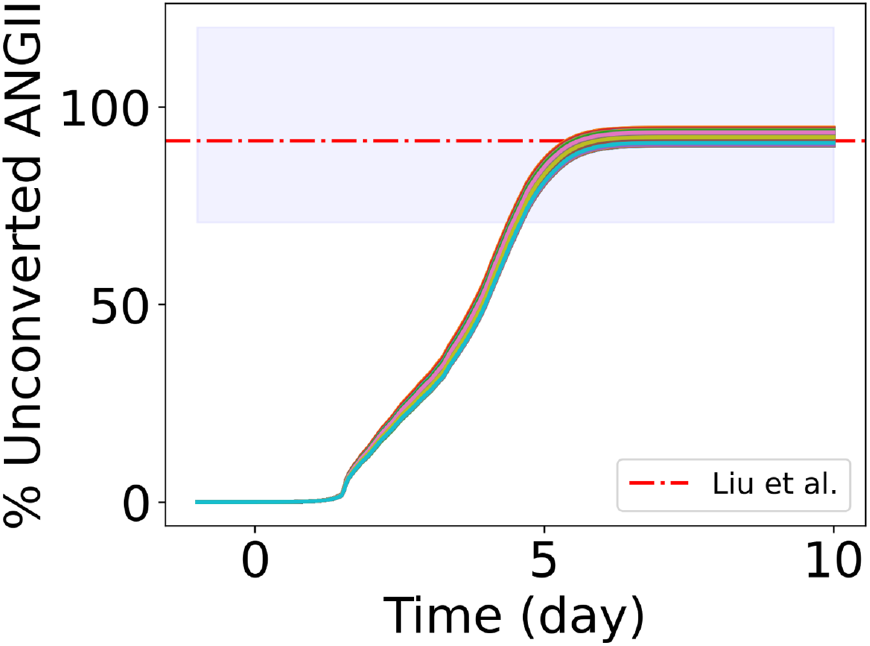
Dynamic profiles of the percentage of unconverted ANGII for 1000 virtual patients with varying initial values of RAS peptides and *ueACE*2_0_ = 1000. The red dashed line and shaded area represent the experimentally reported increases in ANGII in critically ill patients from the mean ANGII of the normal range [2].

**Fig. 8:**
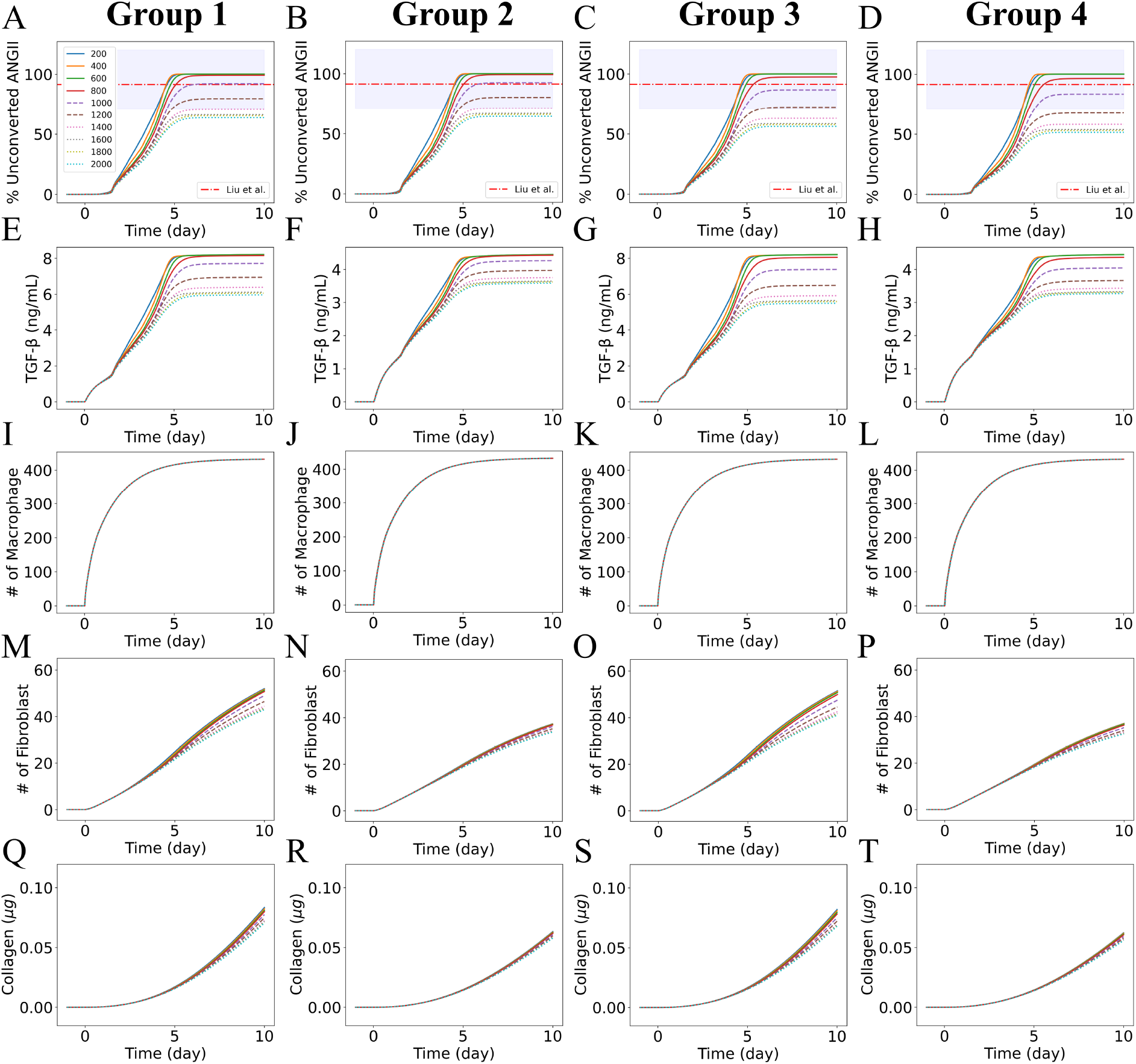
Dynamics of fibrosis model. (Row 1: A–D) unconverted ANGII, (row 2: E–H) TGF-β, (row 3: I–L) macrophages, (row 4: M–P) fibroblasts, and (row 5: O–T) collagen. Patient groups represent each column. Group 1: hypertensive patients with no feedback from ANGII**·**AT1R to renin. Group 2: normotensive patients with no feedback from ANGII**·**AT1R to renin. Group 3: hypertensive patients with feedback from ANGII**·**AT1R to renin. Group 4: normotensive patients with feedback from ANGII**·**AT1R to renin. The red dashed line and shaded area represent the experimentally reported increases in ANGII in critically ill patients from the mean ANGII of the normal range [2]. Note that the common legend for all of the panels appears in the upper left of the figure. The legend shows the age and sex labels for each discrete *ueACE*2_0_ value (see also Table 1). Note that the *y*-axis scales are different for each panel for TGF-β.

Building on the assumption that loss of ACE2 at the tissue scale increases the systemic ANGII, we considered simplified first-order reaction kinetics to account for ANGII-induced TGF-β production from unconverted ANGII (Eq. (14)). The *in vitro* experiments of Lai et al. [66] showed increased TGF-β concentration in human mesangial cells (279–630 pg mL^−1^) with increasing ANGII ranges (0–10^−7^ M). Singh et al. [67] also observed changes in TGF-β concentration in the range 0.827–1.65 ng mL^−1^ with changes of ANGII from the control condition to 10^−8^ M in their *in vitro* experiments on human mesangial cells. The changes in ANGII in our model occur at a much lower range compared to these experiments. However, similar experiments in lung cells could be used to parameterize ANGII-induced TGF-β production rate, *k*_*T*_. Here to determine a reasonable *k*_*T*_ value, we varied *k*_*T*_ in the range of 1*×*10^−4^–10*×*10^−4^ ng mL min^−1^ fmol^−1^ for hypertensive patients with no feedback from ANGII·AT1R to renin and *ueACE*2_0_ = 1000 and observed linear changes in immune cells and collagen dynamics with variations in *k*_*T*_ (Fig. S7). Our goal was to keep TGF-β concentration within the range of experimental observations (0–10 ng/mL) where the TGF-β-dependent functions for fibroblasts were defined [23–25]. We selected *k*_*T*_ = 5 × 10^−4^ ng mL min^−1^ fmol^−1^ to simulate the dynamics of immune cells and collagen for different age and sex groups. Fig. 8 shows the dynamics of the fibrosis model species: unconverted ANGII, TGF-β, macrophages, fibroblasts, and collagen. Fig. S8 shows the dose-response of *ueACE*2_0_ to fibrosis model species at ten days post-infection for the four patient groups described in Section 3.4. Our simulated results show increased macrophage and fibroblast populations and increased collagen deposition over time, consistently across all groups and *ueACE*2_0_ values. However, we observed only slight variations in numbers of fibroblasts and collagen deposition with aging in both sexes across initial *ueACE*2_0_ values (Fig. S8). Fig. S9 shows predictions of the collagen area fraction from our earlier COVID-19 fibrosis model [23] that did not include RAS effects. The collagen area fraction varied depending on age and sex (*ueACE*2_0_ values), and the variations increased from day 10 to day 15, beyond the length of time examined in Fig. 8. Additionally, the effects of the statistics of stochastic outcomes from 15 replications of the ABM were apparent in the nonlinear effects, particularly for *ueACE*2_0_ = 600. Also, the larger values had diminishing effects on collagen area fraction or perhaps an upper bound of some sort was being reached with stochastic fluctuation. These are not the main results of the present model, but shown for comparison purpose to our earlier work [23].

### 3.6. Sensitivity analysis

The dynamic local sensitivity analysis for parameters was conducted to quantify the sensitivity of model output ANGII dynamics to input parameters (Eq. (19)). The analysis was performed for the parameters for submodel 2: *β*_0_, *k*_*A*_, *c*_*A*_, *c*_*R*_, *k*_*ACE*2**·***ANGII*_, *k*_*ACE*2_, *c*_*AP A*_, *c*_*AT* 1_, and *c*_*AT* 2_. We ran the sensitivity analysis using *m*_*S*_ = {0.1, 0.3, 0.5, 0.7, 0.9, 1.1, 1.5, 2, 5, 10} as multiplicative perturbations in the inputs. Figs. 9 and S10 show the local sensitivity for parameters for hypertensive patients with no feedback from ANGII·AT1R to renin (Group 1) and with feedback from ANGII·AT1R to renin (Group 3), all at *ueACE*2_0_ = 1000. The sensitivity analysis of ANGII showed parameters *β*_0_, *c*_*A*_, and *k*_*ACE*2**·***ANGII*_ changed the steady state of ANGII before infection in Group 1 and remained sensitive after infection. *β*_0_ is the renin production rate, *c*_*A*_ is the ANGI to ANGII conversion rate, and *k*_*ACE*2**·***ANGII*_ is the ANGII and ACE2 receptor binding rate. With the feedback in Group 3, two additional parameters were sensitive: *c*_*AT* 1_, the ANGII and type 1 receptor binding rate, and *c*_*R*_, the plasma renin activity on AGT. Before infection *β*_0_ and *c*_*A*_ modulated ANGII response positively, and *k*_*ACE*2**·***ANGII*_ and *c*_*AT* 1_ modulated ANGII response negatively. After the infection, *c*_*A*_, *k*_*ACE*2**·***ANGII*_, and *c*_*R*_ modulated ANGII response positively. We observed a constant sensitivity of *β*_0_ for Group 1 and a decreasing sensitivity of *β*_0_ after an initial increase for Group 3 (Fig. 9). The sensitivity index of *β*_0_ at day 10 for Group 1 was 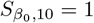, whereas Group 3 range was 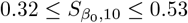 with increasing *m*_*S*_ (Fig. S10). The parameter *c*_*R*_ is sensitive for Group 3 after infection, and the range of sensitivity was 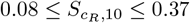 with decreasing *m*_*S*_ (Fig. S10). *c*_*AT*1_ is also sensitive for Group 3, and the sensitivity increases in the negative direction with decreasing *m*_*S*_ (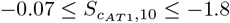, Fig. S10). We also observed a slight negative sensitivity of parameters *c*_*AP A*_ and *c*_*AT*2_ for both Groups 1 and 3.

**Fig. 9:**
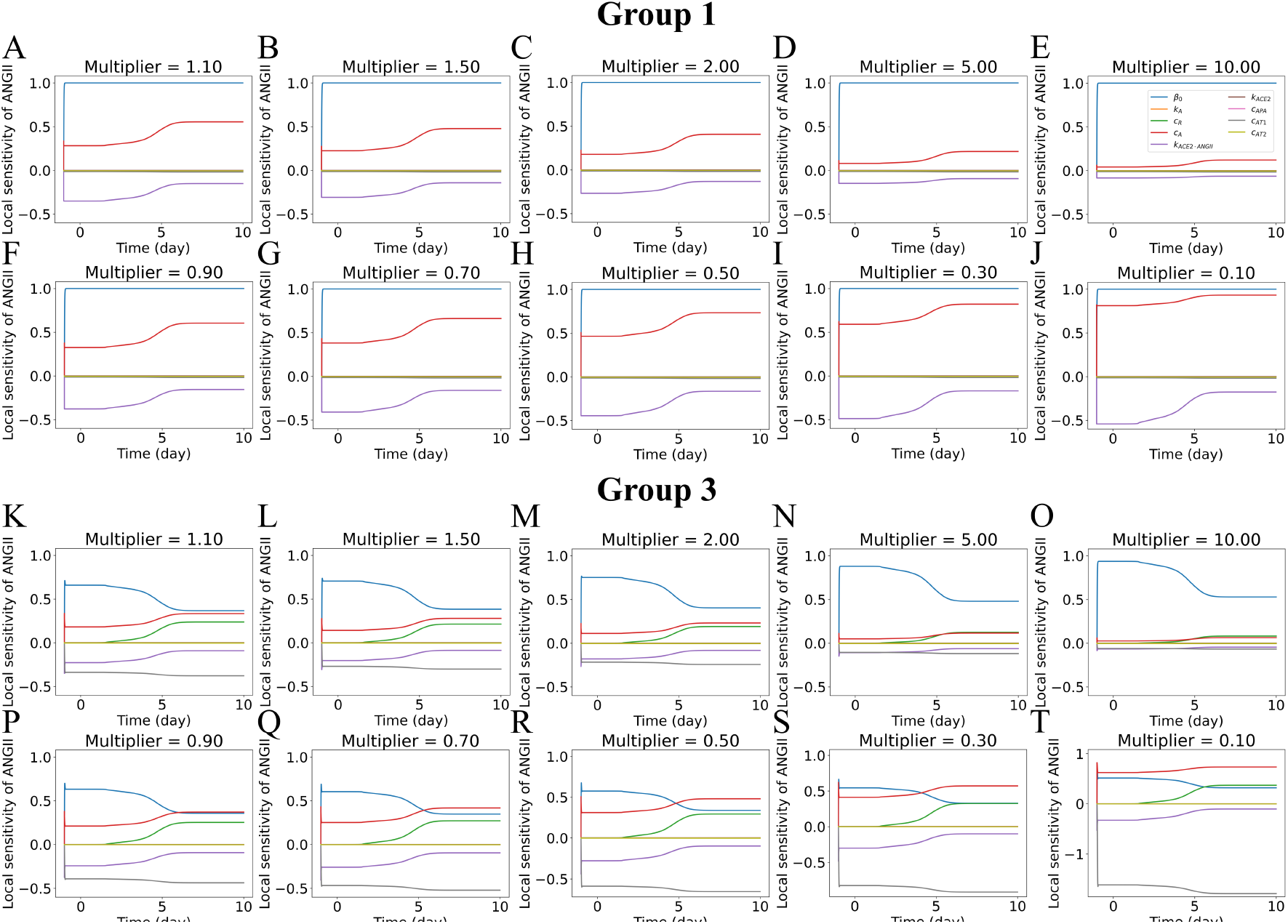
Dynamic local sensitivity analysis of parameters for ANGII for hypertensive patients and *ueACE*2_0_ = 1000. (A–J) Group 1 with no feedback from ANGII**·**AT1R to renin. (K–T) Group 3 with feedback from ANGII**·**AT1R to renin. The parameters are scaled by a multiplier (*m*_*S*_). The values of *m*_*S*_ increase across the first and third rows: (A, K) 1.1, (B, L) 1.5, (C, M) 2, (D, N) 5, (E, O) 10. Values of *m*_*S*_ decrease across the second and fourth rows: (F, P) 0.9, (G, Q) 0.7, (H, R) 0.5, (I, S) 0.3, and (J, T) 0.1. Note that the common legend for all of the panels appears in the upper right of the figure and that the *y*-axis scales are different for the last three panels for Group 3 to accommodate changes in *c*_*AT*1_.

### 3.7. Limitations

In the current model, we used discrete *ueACE*2_0_ values for age and sex groups. However, a continuous *ueACE*2_0_ range may accurately predict the effects of age and sex. We also considered complete destruction of tissue for *ueACE*2_0_ ≤ 600, which might change depending on the patient’s premorbid conditions. Our earlier COVID-19 lung tissue fibrosis model [23] identified M2 macrophages as the key regulators for fibrosis. In this model, we only used experimental data of TGF-β-dependent macrophage recruitment for up to 1 ng mL^−1^ and assumed a constant response from 1–10 ng mL^−1^ (Fig. S1). So, there is a need for experimental studies on TGF-β-dependent macrophage recruitment for TGF-β ranges of 1–10 ng mL^−1^ to remove this assumption in our model, which may enhance the effects of patient differences in the fibrosis model. Although the current model predicted the variations in RAS peptides depending on age and sex, the effect of patient differences is modest for the dynamics of collagen (submodel 3). However, we observed variations in collagen area fraction depending on age and sex in our COVID-19 lung tissue model (submodel 1). In the current workflow, we run the COVID-19 lung tissue model and RAS model sequentially. Instead running them simultaneously would couple the additional systemic influx of immune cells due to dysregulation in RAS and may better simulate the patient differences in collagen deposition.

## 4. Conclusions

Heterogeneity in the severity of COVID-19 disease depends on the patient-specific premorbid conditions, age, and sex differences. We developed an integrated mathematical model to investigate and quantify the effects of these heterogeneous factors on RAS during COVID-19. We identified viral-infection-induced cell death as a major reduction source of ACE2 and did not observe any significant difference due to viral-bound ACE2. Our simulated results showed variations in *ueACE*2_0_ due to age and sex are significant determinants in the dynamics of RAS during COVID-19.

We observed increased disease severity with aging and significant variations between male and female patients in the older and middle-aged groups. Our *in silico* results predicted outcomes for the hypothesized mechanisms were able to explain conflicting RAS peptide alterations in ANGI and ANGII from two previous experimental studies of patients with different COVID-19 severity [1, 6] by considering the variations in homeostasis RAS peptides due to premorbidity and feedback of ANGII·AT1R to renin. The loss of ACE and ACE2 via the death of lung cells resulted in a reduction of RAS peptides. However, the increase in systemic ANGII may result from the loss of ACE2 in the lung tissue. The model also identified that variations in the homeostasis concentrations of RAS peptides due to premorbidity and feedback signaling from ANGII·AT1R to renin are important factors in the patient-specific variations in RAS. We predicted systemic immune recruitment and collagen deposition due to RAS alteration during COVID-19. The model can be calibrated with patient-specific RAS peptides and enzyme concentration, fibrotic mediators from bleomycin-induced tissue fibrosis model, and autopsy and biopsy tissue samples to evaluate detailed dynamics of RAS and fibrosis pathways to develop personalized treatments.

## 5. Acknowledgments

This work was supported by the National Institutes of Health grant R35 GM133763 and the University at Buffalo. We also acknowledge support provided by the Center for Computational Research at the University at Buffalo http://hdl.handle.net/10477/79221. We thank the scientific community for model feedback, including Paul Macklin, Michael Getz, Amber M. Smith, Thomas Hillen, and Yafei Wang.

## 6. Supplementary Material

Supplementary Material associated with this article can be found in the online version.

