## Supplementary Material for "Mathematical Modeling of Impacts of Patient Differences on Renin-Angiotensin System and Applications to COVID-19 Lung Fibrosis Outcomes"

### S1. Workflow of COVID-19 tissue model

The COVID-19 tissue model has 8 cell types and 91 biological hypotheses (agent-based model rules). We listed all the biological hypotheses and agent-based modeling workflow in the supporting information of our earlier manuscript [1]. Here, we summarize the methodology of the COVID-19 tissue model from our earlier manuscript [1].

The COVID-19 tissue model structure is developed in ABM framework PhysiCell [2], which is coupled to BioFVM [3]. BioFVM solves partial differential equations for chemicals considered to be continuous (as opposed to the discrete agents) with terms for diffusion, decay, uptake, and release of multiple substances simultaneously. The extracellular viral transport, uptake by adhering to cell surface ACE2, and virion export from the cell are modeled using BioFVM formulation with Neumann boundary conditions:

$$\frac{\partial \rho_v}{\partial t} = D_v \nabla^2 \rho_v + \sum_{cells\ i} \delta(x - x_i)(-U_i V_i \rho_v + E_i) \tag{S1}$$

where $\rho_v$ is the concentration or population density of the viruses, $D_v$ is the diffusion coefficient, $U_i$ is the virion uptake rate by the cell, $V_i$ is the cell volume, and $E_i$ is the virion export rate from the cell. Here, $D_v = 2.5$ µm$^2$min$^{-1}$ [4, 5].

Virions bind with unoccupied external ACE2 (ueACE2) using a continuum to discrete transition rule based on the binding flux:

$$dv_t = r_b V_c(t) v_e(t) ueACE2(t) \tag{S2}$$

where $r_b$ is the binding rate between virion and ueACE2, $V_c(t)$ is the volume of the cell, $v_e(t)$ is the density of the virions near a cell, and $ueACE2(t)$ is the number of ueACE2 at time $t$. The continuum to discrete transition rule allowed all nearby whole virions uptake when $dv_t > 1$. For fractional virion or for $dv_t < 1$, which resulted from the continuum rule in Eq S1, virion uptake depends on the probabilistic rule. The fractional virion uptake occurs when a random number drawn from a uniform distribution is greater than the fractional binding flux ($U(0, 1) > dv_t$, where $0 < dv_t < 1$).

The receptor kinetics and intracellular viral replication kinetics are modeled as a set of ordinary differential equations (ODEs), and corresponding parameters are selected from the experimental characterization of SARS-CoV entry into host cells [6]. The assembled virions also initiate cell death response, which is modeled as Hill function to relate the apoptosis rate of a cell to assembled virions. The infected cells secrete chemokine until the cell dies through lysis or CD8+ T cell-induced apoptosis. Details of the intracellular virus model for replication kinetics, viral response, and receptor trafficking are described in much greater detail in the earlier model manuscript [7].

The resident and newly recruited macrophages move along the gradients of chemokine and debris, phagocytose dead cells, and secrete pro-inflammatory cytokines. We consider five states of macrophages: inactivated, M1 phenotype, M2 phenotype, hyperactive, and exhausted. Both M1 and M2 phenotypes can also exist in hyperactive and

exhausted states. Neutrophils are recruited into the tissue by pro-inflammatory cytokines, move along the gradients of chemokine and debris, phagocytose dead cells, and uptake virions. The resident DCs chemotaxis along the gradients of chemokines, activated by infected cells and viruses, and a portion of the activated DCs egress out of the tissue to the lymph node to induce activation of virus-specific CD4+ and CD8+ T cells. The activated T cells are recruited in the tissue from the lymph node. CD8+ cells attempt to attach with infected cells based on PhysiCell's mechanical interaction and cumulative attachment above a threshold time (25 min) causes apoptosis of the infected cells. The contact between CD4+ T cells and macrophages induces a hyperactive state in macrophages, enabling them to phagocytose live infected cells. The out-of-bound cells from the simulated domain are pushed back into the tissue section by changing the direction away from the edge.

Extracellular densities of chemokines and pro-inflammatory cytokines are modeled using standard BioFVM formulation with Neumann boundary conditions:

$$\frac{\partial \rho}{\partial t} = D\nabla^2 \rho_v - \lambda\rho + \sum_{cells\ i} \delta(x - x_i)(S_i(\rho_i^* - \rho) - U_i\rho)V_i \tag{S3}$$

where $D$ is the diffusion coefficient, $\lambda$ is the decay rate, $S_i$ is the secretion rate, $U_i$ is the uptake rate, and $\rho_i^*$ is the saturation density. For chemokines, pro-inflammatory cytokines, and debris, diffusion coefficients are assumed to be the same as the diffusion coefficient for monoclonal antibodies [8] and set at 555.56 $\mu m^2 min^{-1}$; decay rates are assumed to be the same as the decay of IL-6 [9] and set at $1.02 \times 10^{-2}$ $min^{-1}$. Further details of parameters for immune cells, equations for chemotaxis and recruitment, rules for phagocytosis and CD8+ T cell-induced apoptosis, and proliferation, activation, and clearance of the T cells in the lymph nodes using ODEs are available elsewhere [7, 10, 11].

The death of an infected epithelial cell activates the latent TGF-β embedded in the tissue ECM. The activation of latent stores of TGF-β is represented by creating a stationary secreting agent at the site and time of epithelial cell death. The secreting agents secrete TGF-β for a certain period as the amount of TGF-β stored in the ECM is limited; a death rate of secreting agents ($AT$) is used to represent terminating the activation of latent stores of TGF-β. Here, death rate $AT$ is analogous to the extinction rate of the ECM-bound TGF-β sources and is inversely proportional to the mean duration of those sources.

Macrophages are the mobile sources of TGF-β in this fibrosis model. M1 macrophages are recruited in the initial phase, and CD8+ T cells are recruited in the later stage of infection. Usually, regulatory T cells (Tregs), a subset of CD4+ T cells, are involved in the phenotypic transformation of M1 to M2 macrophages [12]. Although we do not have Tregs in the current model, CD8+ T cells exhibit similar behavior by inhibiting the secretion of pro-inflammatory cytokines from M1 macrophages in the later phase of infection [10]. Since the fibroproliferative phase of DAD occurs in the later phase of the disease, a conditional rule is added in the fibrosis model to follow this pathology: if a CD8+ T cell and an M1 macrophage are within close proximity, then the M1 macrophage stops secreting pro-inflammatory cytokines, transforms to an M2 macrophage, and starts secreting TGF-β. The interaction distance for this transition is calculated using a multiple ($\epsilon$) of the radii of the two interacting immune cells [10]. The transformation of the M1 to M2 phenotype is considered to occur instantaneously when the distance between an M1 macrophage and a CD8+ T cell is $\leq \epsilon$ multiplied by the sum of the radii of the M1 macrophage and the CD8+ T cell. The cell radii are computed in PhysiCell from cell volumes over time as the cell volume changes with the progression of the cell cycle. The initial volume of macrophages is 4849 $\mu m^3$ and that for CD8+ T cell is 478 $\mu m^3$ with nuclear volumes of 10% of the cell volumes; these are the same values specified in the overall model [7].

We simulate extracellular concentrations of TGF-β ($T_\beta$) using the standard BioFVM formulation with Neumann boundary conditions:

$$\frac{\partial T_\beta}{\partial t} = D_{T\beta}\nabla^2 T_\beta - \mu_{T\beta} T_\beta + \sum_{cells\ i} \delta(x - x_i) E_{T\beta,i} \tag{S4}$$

where $t$ is time, $D_{T\beta}$ is the diffusion coefficient of TGF-β, $\mu_{T\beta}$ is the net decay and bulk removal rate, $\delta(x)$ is the discrete Dirac delta function, $x$ is the center of the voxel, $x_i$ is the position of the center of discrete cell $i$, and $E_{T\beta,i}$ is the net export rate of TGF-β from cell $i$. Net export rate $E_{T\beta,i}$ denote the activation rate of latent stores of TGF-β at the damaged site, which we term damaged-site secretion ($DS$), or TGF-β secretion and activation rate from macrophages, which we term macrophage secretion ($MS$).

Initial fibroblasts remain in an inactive state. To maintain the homeostatic population of fibroblasts, we assume that the initial inactive fibroblasts move randomly throughout the domain and do not undergo apoptosis. TGF-β activates the resident inactive fibroblasts. We assume that inactive fibroblasts become active fibroblasts in the presence of TGF-β ($T_\beta > 0$) and switch back to the inactive state in the absence of TGF-β ($T_\beta = 0$). The active fibroblasts and inactive fibroblasts from the active state undergo apoptosis naturally and become dead cells. Both inactive and active fibroblasts are subject to the cycle cell model in PhysiCell where their cell volumes are over updated time as the cell cycle progresses.

The activated fibroblasts chemotax up the gradient of TGF-β. The chemotaxis velocity of a fibroblast ($\vec{v}_{mot}$) depends on the TGF-β concentration in its neighboring regions, fibroblast speed ($s_{mot}$), migration bias ($b$), and time for persistence in a specific trajectory ($\Delta t_{mot}$) [10]:

$$\vec{v}_{mot} = s_{mot} \frac{b\vec{b} + (1-b)\vec{\xi}}{\left\| b\vec{b} + (1-b)\vec{\xi} \right\|} \tag{S5}$$

where $s_{mot}$ is the speed of chemotaxis, $0 \leq b \leq 1$ is the level of migration bias (i.e., $b = 0$ represents no influence of chemotaxis and only random cell migration), $\vec{\xi}$ is a random unit vector direction, and $\vec{b}$ is the migration bias direction. Fibroblasts also persist on their given trajectory for $\Delta t_{mot}$ before a new trajectory is computed. For fibroblasts, $\vec{b}$ is estimated from the gradient of TGF-β concentrations in the neighboring regions [10]:

$$\vec{b} = \frac{\nabla T_\beta}{\|\nabla T_\beta\|} \tag{S6}$$

Chemotaxis of other immune cells along the gradients of pro-inflammatory cytokines, chemokines, and debris in the overall model follows this formulation[10].

New fibroblasts are recruited into the tissue by TGF-β. The number of fibroblasts recruited to the tissue ($N_r$) is determined by integrating the recruitment signal over the whole domain:

$$N_r = r_r \int_\Omega \min\left(1, \max\left(0, \frac{s_{cytokine} - s_{min}}{s_{sat} - s_{min}}\right)\right) dV \Delta t_r \tag{S7}$$

where $\Omega$ is the computational domain, $r_r$ is the recruitment rate (per volume), $s_{min}$ is the minimum recruitment signal, $s_{sat}$ is the saturating or maximum signal, and $s_{cytokine}$ is the recruitment signal that depends on the cytokine concentration. The volume of each grid voxel is $dV = 20 \times 20 \times 20$ $\mu$m$^3$, and the time interval for recruitment of fibroblasts is $\Delta t_r = 10$ min, the same as in the overall model [10]. Eq S7 gives the number of fibroblasts recruited between $t$ and $t + \Delta t_r$. Here, $s_{cytokine} = F_g(T_\beta)$ for the recruitment signal of fibroblasts via the cytokine TGF-β. Recruitment of any immune cell by a specific cytokine in the overall model follows this formulation; proinflammatory cytokine-dependent recruitment of neutrophils and macrophages is already included in the earlier model [10].

The function for TGF-β-dependent recruitment of fibroblasts ($F_g(T_\beta)$)—particularly its specific parameters—comes from previously published works [13, 14]. We selected a threshold value to set the upper range of TGF-β concentration beyond which $F_g(T_\beta)$ is assumed to be constant because of the polynomial nature of the recruitment signal.

$$F_g(T_\beta) = \begin{cases} 0.0492T_\beta^3 - 0.9868T_\beta^2 + 6.5408T_\beta + 7.1092, & \text{if } 0 < T_\beta \leq 10 \\ 22.928, & \text{else if } T_\beta > 10 \\ 0, & \text{otherwise} \end{cases} \tag{S8}$$

The activated fibroblasts deposit collagen continuously. The newly deposited collagen is assumed to not diffuse or degrade. TGF-β-dependent collagen deposition from fibroblasts is described by

$$\frac{\partial C}{\partial t} = \sum_{cells\ i} \delta(x - x_i) E_{C,i} \tag{S9}$$

and

$$E_{C,i} = k_{FC} F_c(T_\beta) = k_{FC} \frac{V_{T\beta} T_\beta}{k_{T\beta} + T_\beta} \tag{S10}$$

where $C$ is the concentration of collagen, $E_{C,i}$ is the net export rate of collagen from fibroblast cell $i$, and $k_{FC}$ is the collagen production rate by fibroblasts. $F_c(T_\beta)$ defines the TGF-β dependence of fibroblast-mediated collagen deposition, which is assumed to follow a Michaelis-Menten form, where $V_{T\beta}$ and $k_{T\beta}$ are the Michaelis-Menten limiting rate and half saturation parameters, respectively.

Details of the parameter estimation of the COVID-19 tissue model is in our earlier manuscript [1]. The simulation update time for the microenvironment using BioFVM formulation is 0.01 min, cell mechanics is 0.1 min, cell processes is 6 min, and cell recruitment is 10 min. The total simulation time is 21,600 min (15 days), and the data output for saving every 60 min.

### S2. Supplementary equations for RAS model

We solved Eqs. (2)–(12) at steady-state represented by Eqs. (S11)–(S20) assuming $c_{mas} = c_{AT2}$ to calculate the patient-specific parameters. The bold symbols denote the unknown in the equations.

$$\boldsymbol{\beta_0} - \frac{\ln 2}{h_R}[\boldsymbol{R}]_{\boldsymbol{0}} = 0 \tag{S11}$$

$$\boldsymbol{k_A} - c_R[\boldsymbol{R}]_{\boldsymbol{0}} - \frac{\ln 2}{h_A}[AGT]_0 = 0 \tag{S12}$$

$$c_R[\boldsymbol{R}]_{\boldsymbol{0}} - \boldsymbol{c_A}[ANGI]_0 ACE_0 - \frac{\ln 2}{h_{A1}}[ANGI]_0 = 0 \tag{S13}$$

$$\begin{aligned}\boldsymbol{c_A}[ANGI]_0 ACE_0 - \boldsymbol{c_{APA}}[ANGII]_0 - \boldsymbol{c_{AT1}}[ANGII]_0 - \boldsymbol{c_{AT12}}[ANGII]_0 \\ - \boldsymbol{k_{ACE2 \cdot ANGII}}[ANGII]_0 ACE2_0 - \frac{\ln 2}{h_{A2}}[ANGII]_0 = 0\end{aligned} \tag{S14}$$

$$\boldsymbol{k_{ACE2}}[ACE2 \cdot ANGII]_0 - c_{mas}[ANG1\text{–}7]_0 - \frac{\ln 2}{h_{A17}}[ANG1\text{–}7]_0 = 0 \tag{S15}$$

$$\boldsymbol{k_{ACE2 \cdot ANGII}}[ANGII]_0 ACE2_0 - \boldsymbol{k_{ACE2}}[ACE2 \cdot ANGII]_0 = 0 \tag{S16}$$

$$\boldsymbol{c_{APA}}[ANGII]_0 - \frac{\ln 2}{h_{A4}}[ANGIV]_0 = 0 \tag{S17}$$

$$\boldsymbol{c_{AT1}}[ANGII]_0 - \frac{\ln 2}{h_{AT1}}[ANGII \cdot AT1R]_0 = 0 \tag{S18}$$

$$\boldsymbol{c_{AT2}}[ANGII]_0 - \frac{\ln 2}{h_{AT2}}[ANGII \cdot AT2R]_0 = 0 \tag{S19}$$

$$c_{mas}[ANG1\text{–}7]_0 - \frac{\ln 2}{h_{mas}}[\boldsymbol{MAS \cdot ANG1\text{–}7}]_{\boldsymbol{0}} = 0 \tag{S20}$$

$$c_{mas} = \boldsymbol{c_{AT2}} \tag{S21}$$

### S3. Supplementary figures

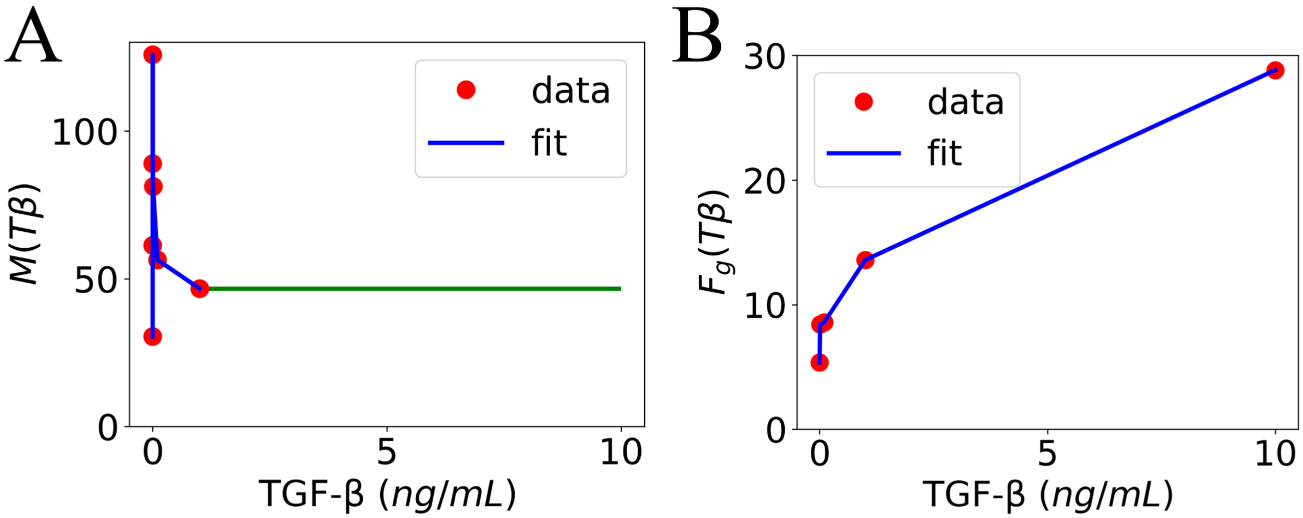


Fig. S1: TGF-$\beta$ dependent recruitment of (A) macrophages and (B) fibroblasts. The experimental data from Wahl et al. [15] is represented by solid red circles. Blue lines show linear fitting between data points, and the green line extrapolates from the endpoint data.

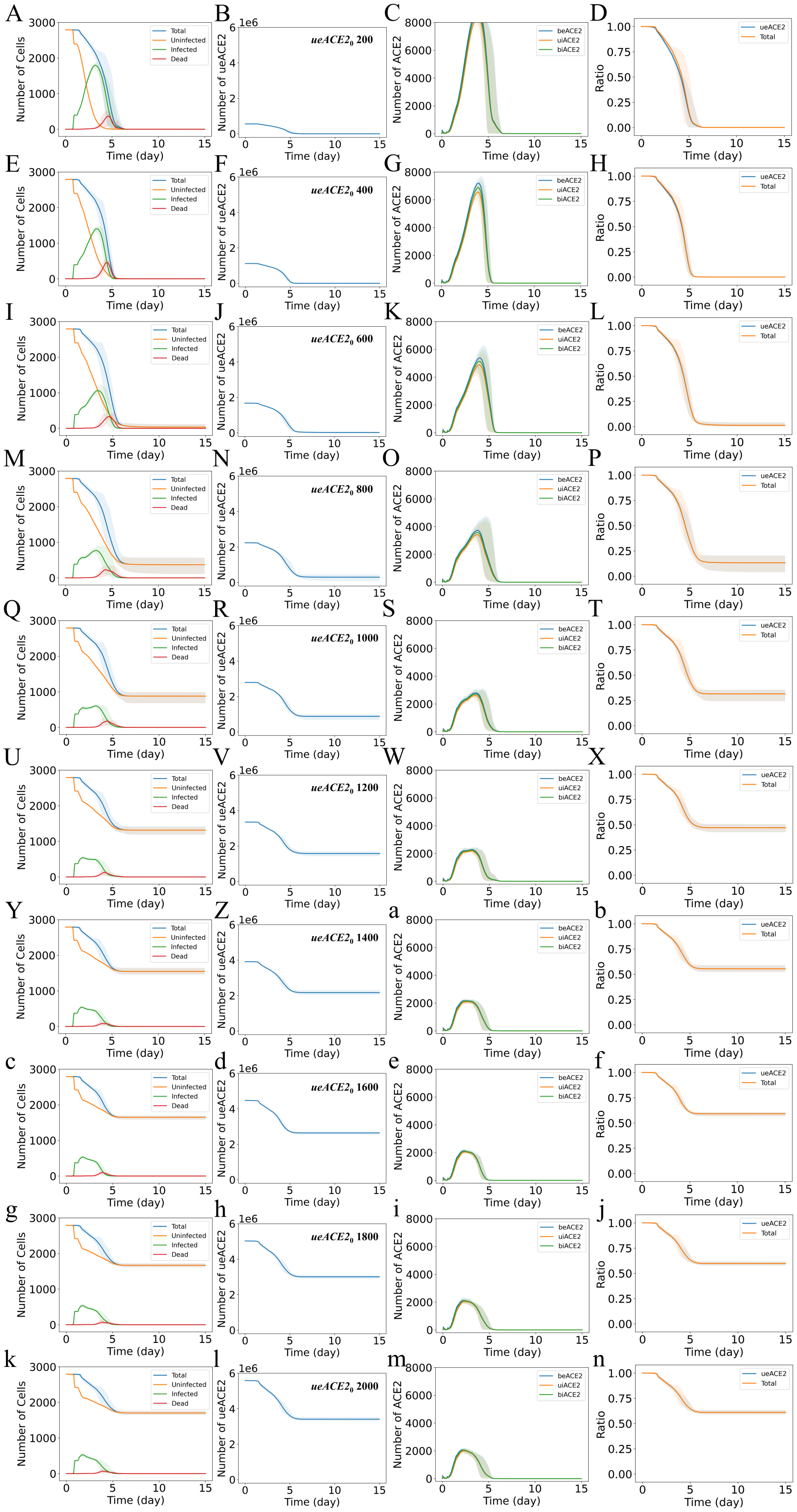


Fig. S2: Epithelial cells and ACE2 response to different initial values of unbound external ACE2 per epithelial cell ($ueACE2_0$). Dynamics of (column 1: A, E, I, M, Q, U, Y, c, g, k) total, uninfected, infected, and dead epithelial cells; (column 2: B, F, J, N, R, V, Z, d, h, l) tissue-wide unbound external ACE2 (ueACE2) (column 3: C, G, K, O, S, W, a, e, i, m) tissue-wide bound external ACE2 (beACE2), unbound internal ACE2 (uiACE2), and bound internal ACE2 (biACE2); and (column 4: D, H, L, P, T, X, b, f, j, n) normalized comparison between the dynamics of tissue-wide ueACE2 and total cells. Each row represents a fixed $ueACE2_0$ value in the range of 200–2000 receptors per cell, as labeled in column 2. The numbers of ueACE2 on the $y$-axes of column 2 plots (B, F, J, N, R, V, Z, d, h, and l) denote the total number of ueACE2 receptors in the virtual lung tissue. The solid curves represent the means, and shaded areas represent the 5th and 95th percentiles of 15 iterations.

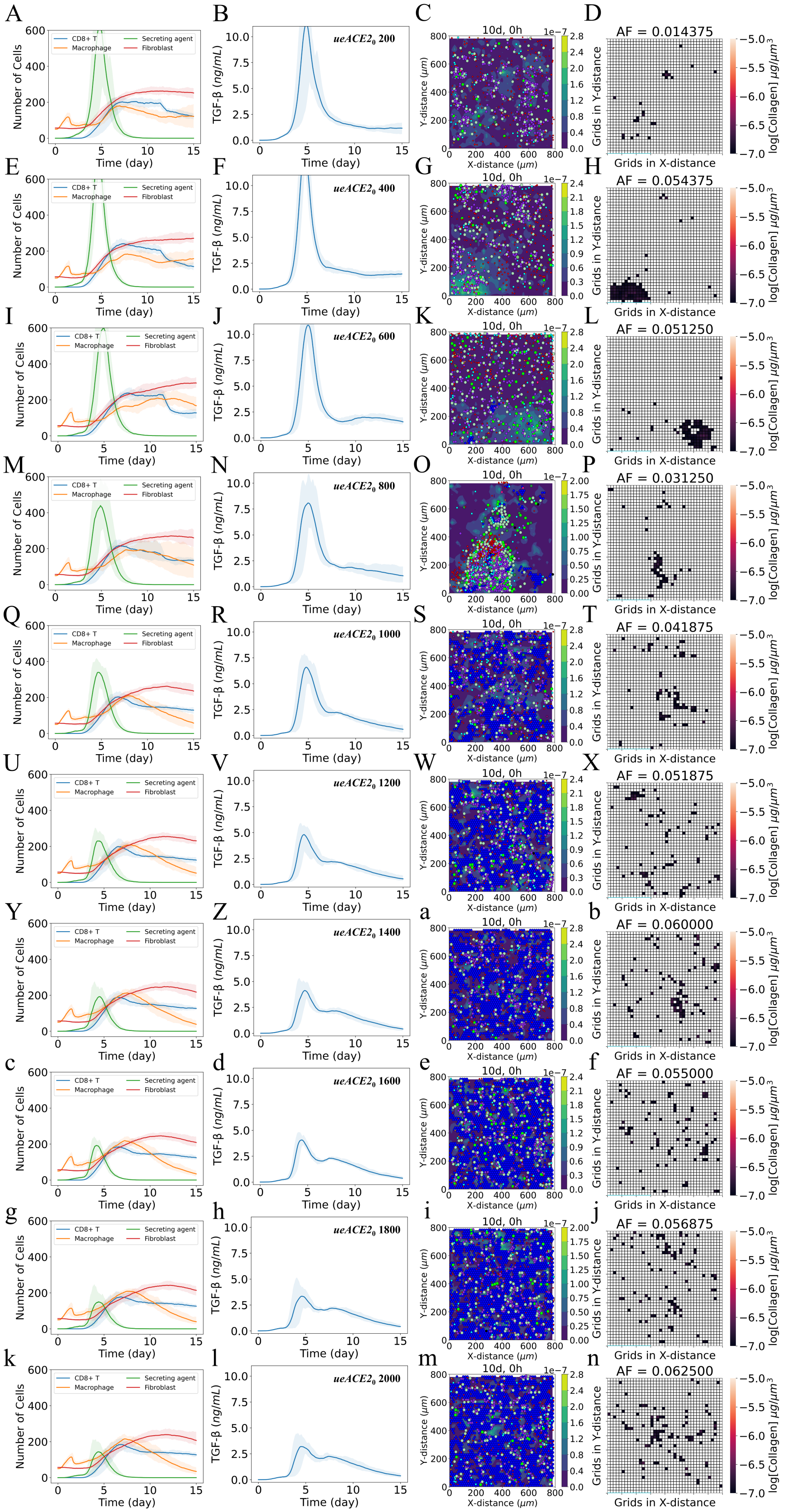


Fig. S3: Virtual lung tissue response to different initial values of unbound external ACE2 per epithelial cell ($ueACE2_0$) and resulting local fibrosis. Dynamics of (column 1: A, E, I, M, Q, U, Y, c, g, k) CD8+ T cells, macrophages, fibroblasts, and the secreting agents that result from damaged epithelial cells; (column 2: B, F, J, N, R, V, Z, d, h, l) mean TGF-$\beta$ concentration; (column 3: C, G, K, O, S, W, a, e, i, m,) collagen deposited (µg $\mu m^{-3}$) at damaged sites in tissue at day 10; and (column 4: D, H, L, P, T, X, b, f, j, n)) heat map showing collagen area fraction (AF) above the threshold value of $1 \times 10^{-7}$ µg $\mu m^{-3}$. Each row represents a fixed $ueACE2_0$ value in the range of 200–2000 receptors per cell, as labeled in column 2. The solid curves represent the means, and shaded areas represent the 5th and 95th percentiles of 15 iterations.

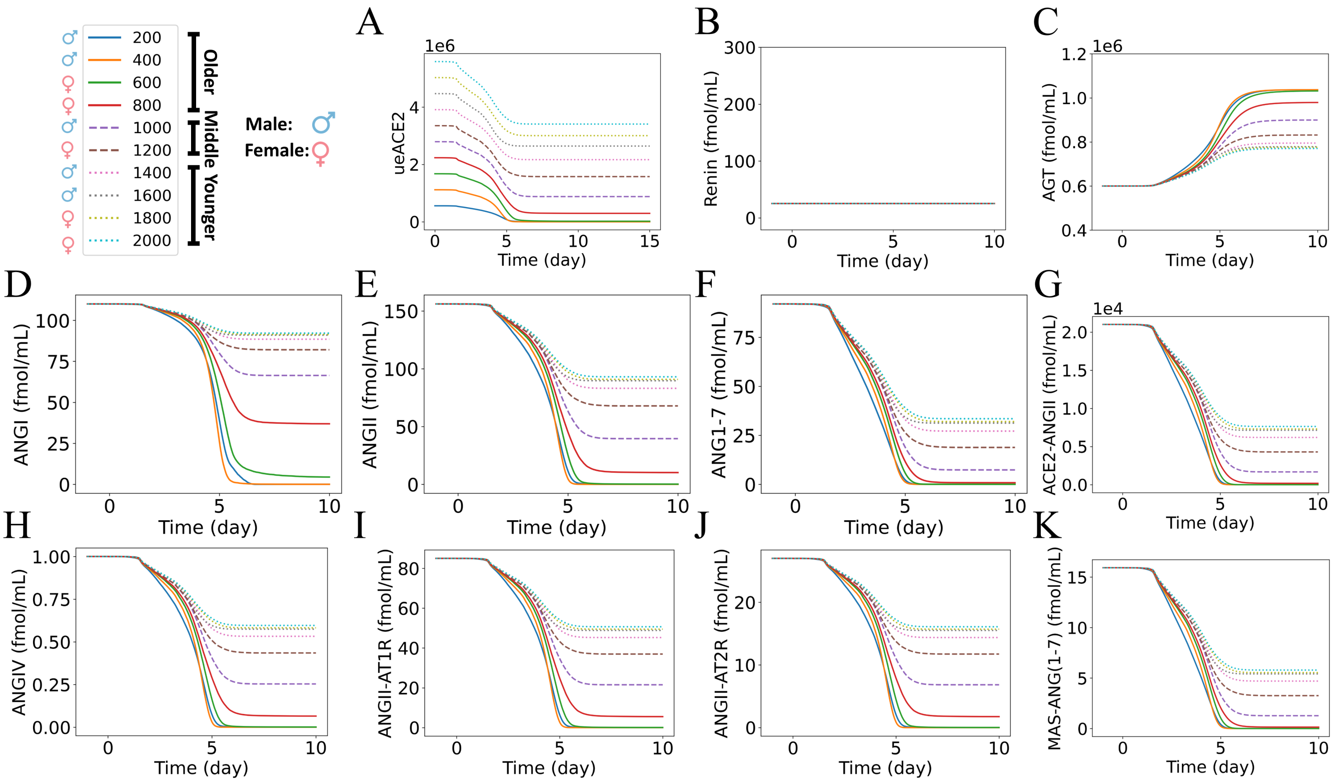


Fig. S4: Dynamics of RAS peptides in response to different initial values of unbound external ACE2 per epithelial cell ($ueACE2_0$) for Group 1: hypertensive patients with no feedback from ANGII·AT1R to renin. Dynamics of (A) tissue-wide unbound external ACE2, (B) renin, (C) AGT, (D) ANGI, (E) ANGII, (F) ANG1–7, (G) ACE2·ANGII, (H) ANGIV, (I) ANGII·AT1R, (J) ANGII·AT2R, and (K) MAS·ANG(1-7) for $ueACE2_0$ values in the range 200–2000 receptors per cell. The $y$-axis of A denotes the total number of ueACE2 receptors in the virtual lung tissue. The legend shows the age and sex labels for each discrete $ueACE2_0$ value.

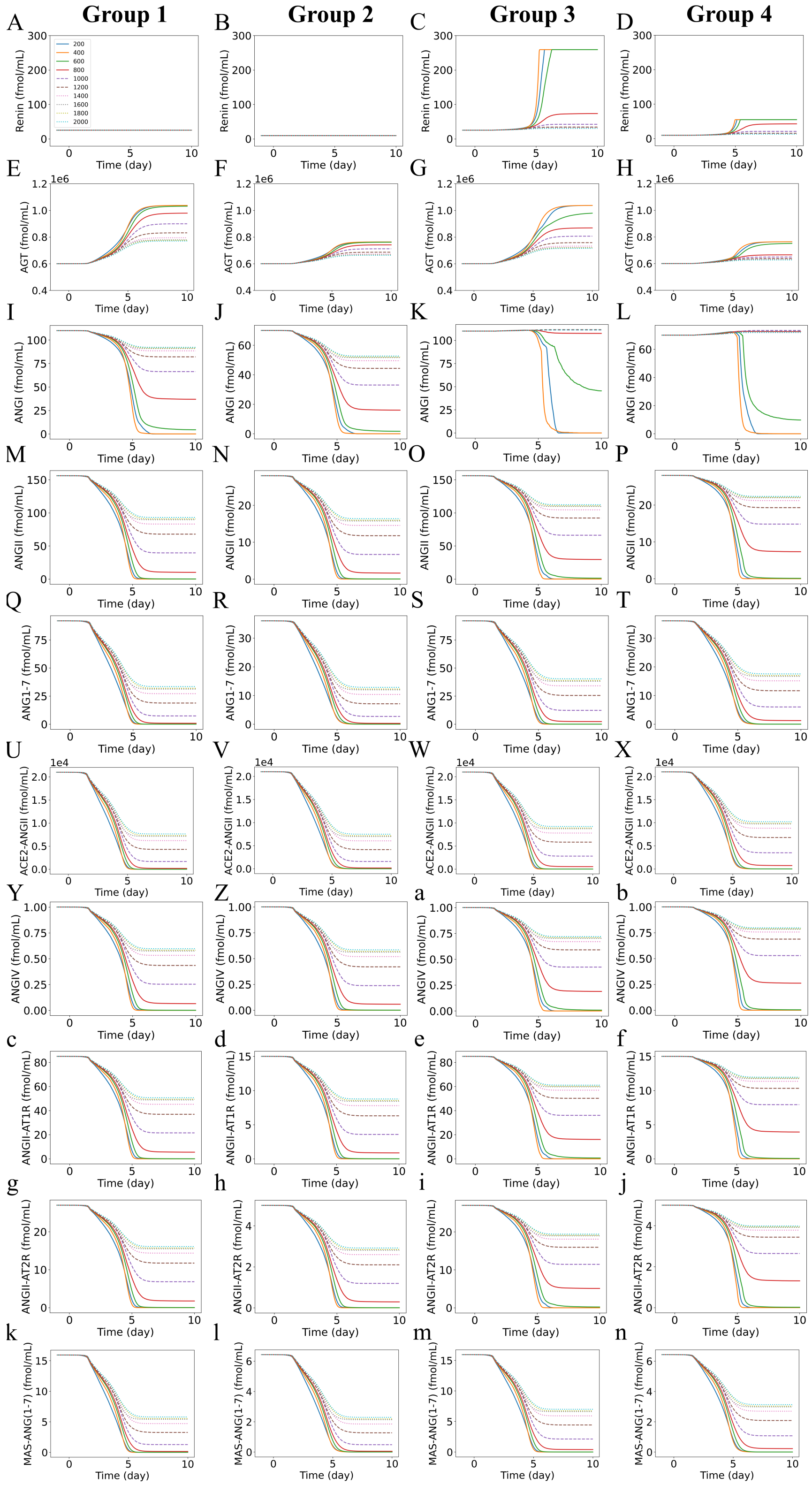


Fig. S5: Dynamics of RAS peptides in response to different initial values of unbound external ACE2 per epithelial cell ($ueACE2_0$) for the four patient groups. Dynamics of (row 1: A–D) renin, (row 2: E–H) AGT, (row 3: I–L) ANGI, (row 4: M–P) ANGII, (row 5: Q–T) ANG1–7, (row 6: U–X) ACE2·ANGII, (row 7: Y–b) ANGIV, (row 8: c–f) ANGII·AT1R, (row 9: g–j) ANGII·AT2R, and (row 10: k–n) MAS·ANG(1-7). Patient groups represent each column. Group 1 (column 1): hypertensive patients with no feedback from ANGII·AT1R to renin. Group 2 (column 2): normotensive patients with no feedback from ANGII·AT1R to renin. Group 3 (column 3): hypertensive patients with feedback from ANGII·AT1R to renin. Group 4 (column 4): normotensive patients with feedback from ANGII·AT1R to renin. Group 1 is repeated from Fig. S4 for comparison to other groups. Note that the common legend for all of the panels appears in the upper left of the figure. The age and sex labels for each discrete $ueACE2_0$ value are shown in the legend of Fig. S4. Note that the $y$-axis scales are different for each subfigure in a comparison row.

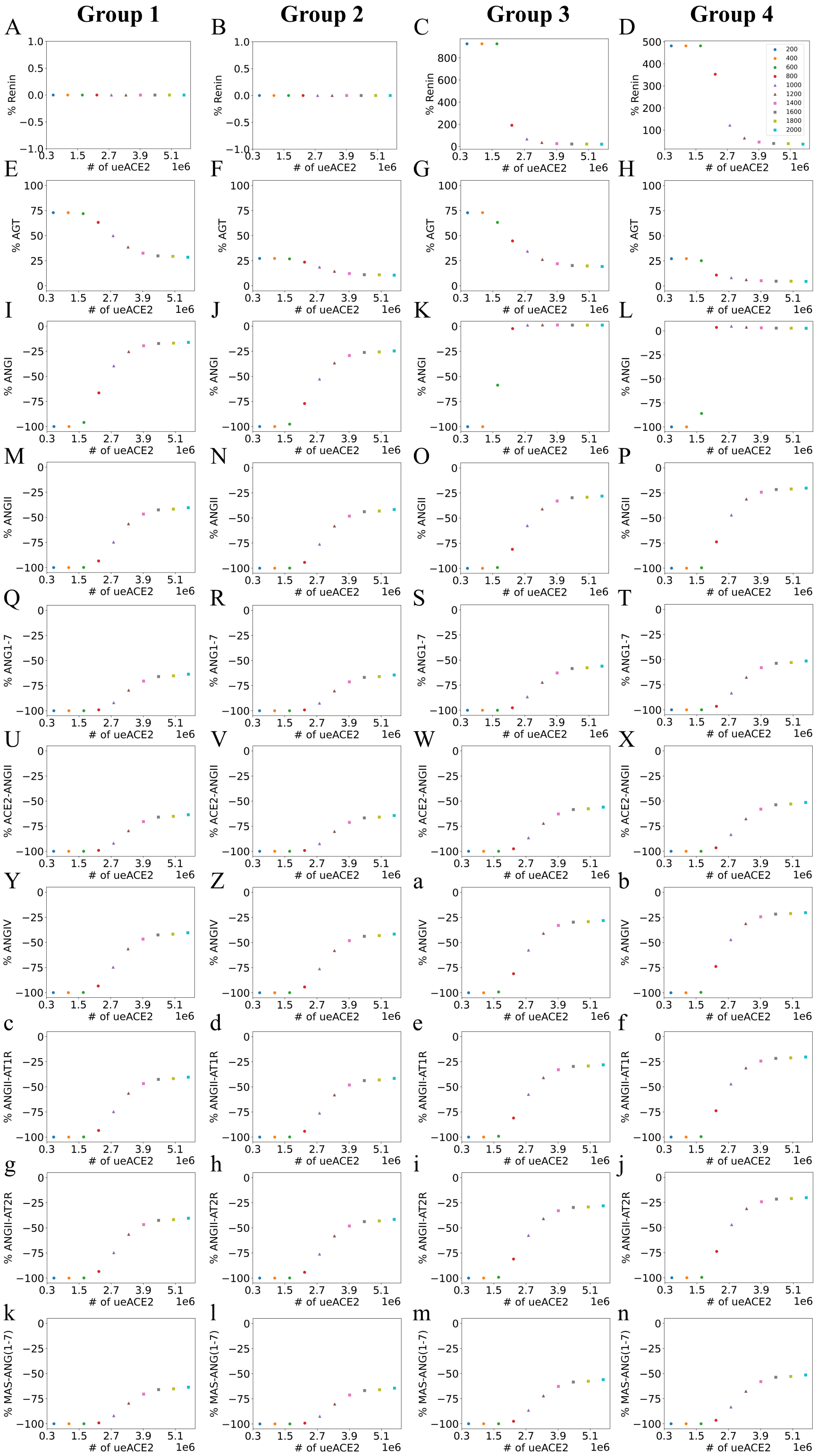


Fig. S6: Dose-response at 10 days after infection to different initial values of unbound external ACE2 per epithelial cell ($ueACE2_0$) for percent change in RAS peptides compared to baseline: (row 1: A–D) renin, (row 2: E–H) AGT, (row 3: I–L) ANGI, (row 4: M–P) ANGII, (row 5: Q–T) ANG1–7, (row 6: U–X) ACE2·ANGII, (row 7: Y–b) ANGIV, (row 8: c–f) ANGII·AT1R, (row 9: g–j) ANGII·AT2R, and (row 10: k–n) MAS·ANG(1-7). Patient groups represent each column. Group 1 (column 1): hypertensive patients with no feedback from ANGII·AT1R to renin. Group 2 (column 2): normotensive patients with no feedback from ANGII·AT1R to renin. Group 3 (column 3): hypertensive patients with feedback from ANGII·AT1R to renin. Group 4 (column 4): normotensive patients with feedback from ANGII·AT1R to renin. Note that the common legend for all of the panels appears in the upper right of the figure. The age and sex labels for each discrete $ueACE2_0$ value are shown in the legend of Fig. S4. Note that the $y$-axis scales are different for each subfigure in row 1.

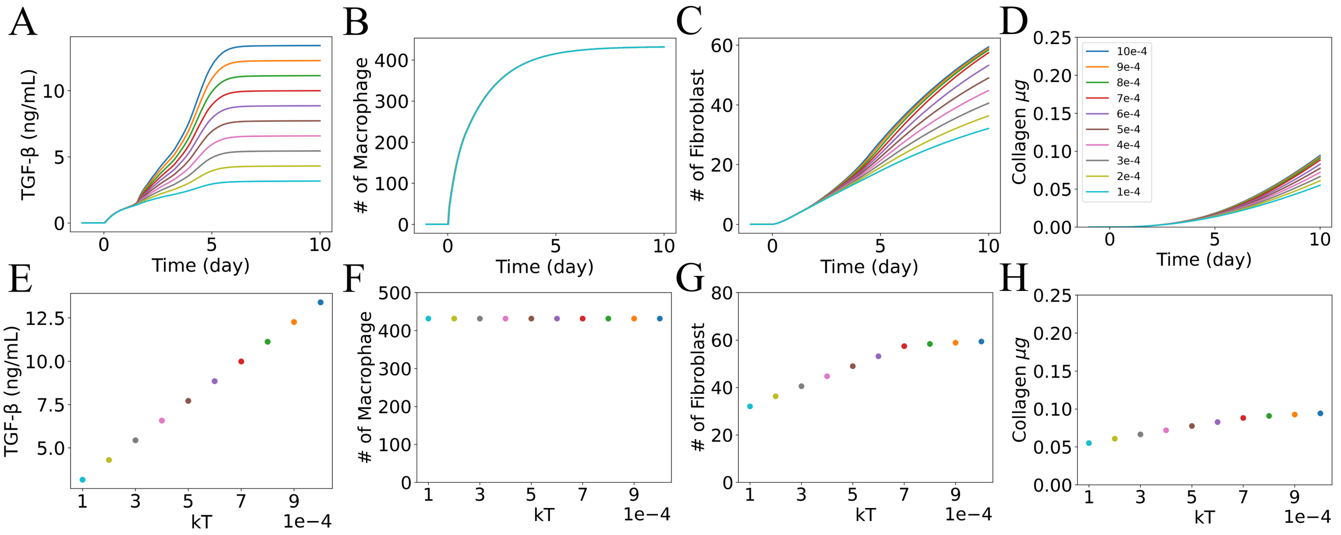


Fig. S7: Effect of ANGII-induced TGF-β production rate ($k_T$) on TGF-$\beta$, immune cells, and collagen deposition for $ueACE2_0 = 1000$ and Group 1: hypertensive patients with no feedback from ANGII·AT1R to renin. Dynamics of (A) TGF-$\beta$, (B) macrophages, (C) fibroblasts, and (D) collagen. Dose-response at 10 days after infection as a function of $k_T$ input for (E) TGF-$\beta$, (F) macrophages, (G) fibroblasts, and (H) collagen. Note that the common legend for all of the panels appears in the upper left of the figure, where the colors correspond to $k_T$ values. Note that the $y$-axis scales are different for each panel.

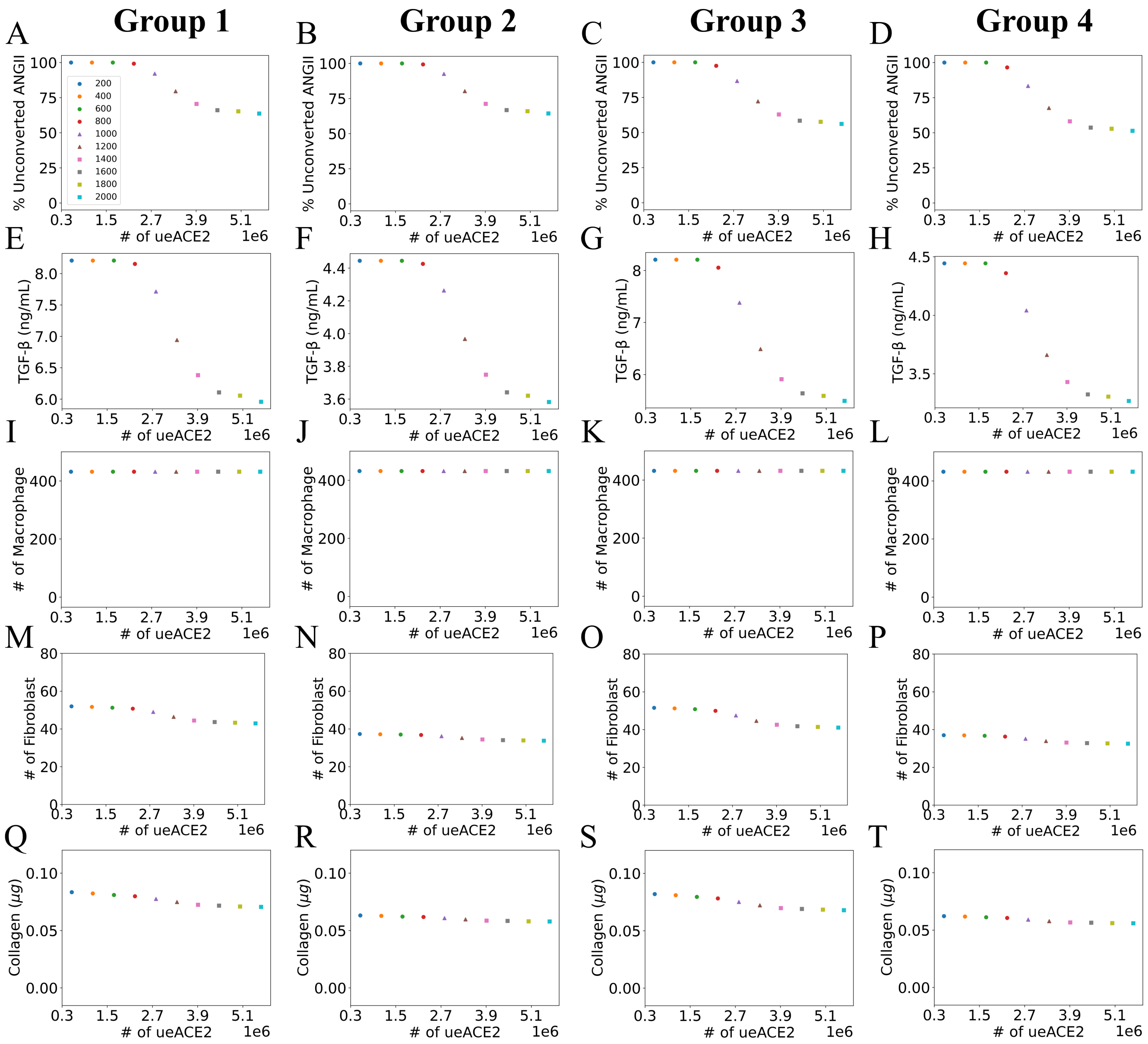


Fig. S8: Dose-response at 10 days after infection to different initial values of unbound external ACE2 per epithelial cell ($ueACE2_0$) for (row 1: A–D) unconverted ANGII, (row 2: E–H) TGF-$\beta$, (row 3: I–L) macrophages, (row 4: M–P) fibroblasts, and (row 5: O–T) collagen. Patient groups represent each column. Group 1 (column 1): hypertensive patients with no feedback from ANGII·AT1R to renin. Group 2 (column 2): normotensive patients with no feedback from ANGII·AT1R to renin. Group 3 (column 3): hypertensive patients with feedback from ANGII·AT1R to renin. Group 4 (column 4): normotensive patients with feedback from ANGII·AT1R to renin. Note that the common legend for all of the panels appears in the upper left of the figure. The age and sex labels for each discrete $ueACE2_0$ value are shown in the legend of Fig. S4.

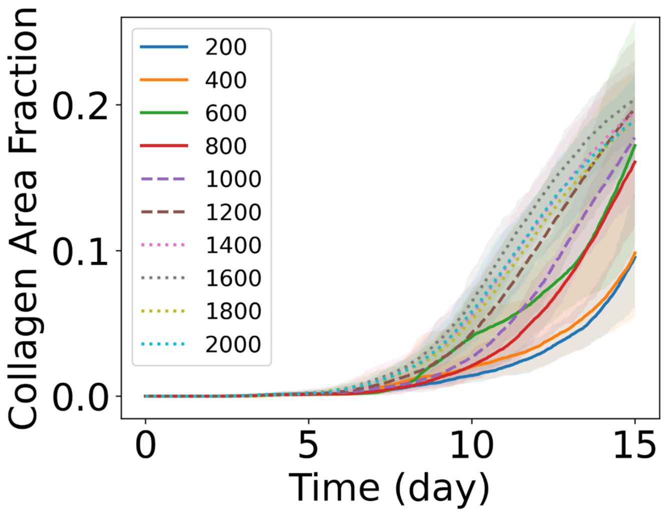


Fig. S9: Dynamics of collagen area fraction in the virtual lung tissue in response to different initial values of unbound external ACE2 per epithelial cell ($ueACE2_0$). The solid curves represent the mean of predictions, and shaded areas represent the predictions between the 5th and 95th percentile of 15 replications of the agent-based model. The age and sex labels for each discrete $ueACE2_0$ value are shown in the legend of Fig. S4.

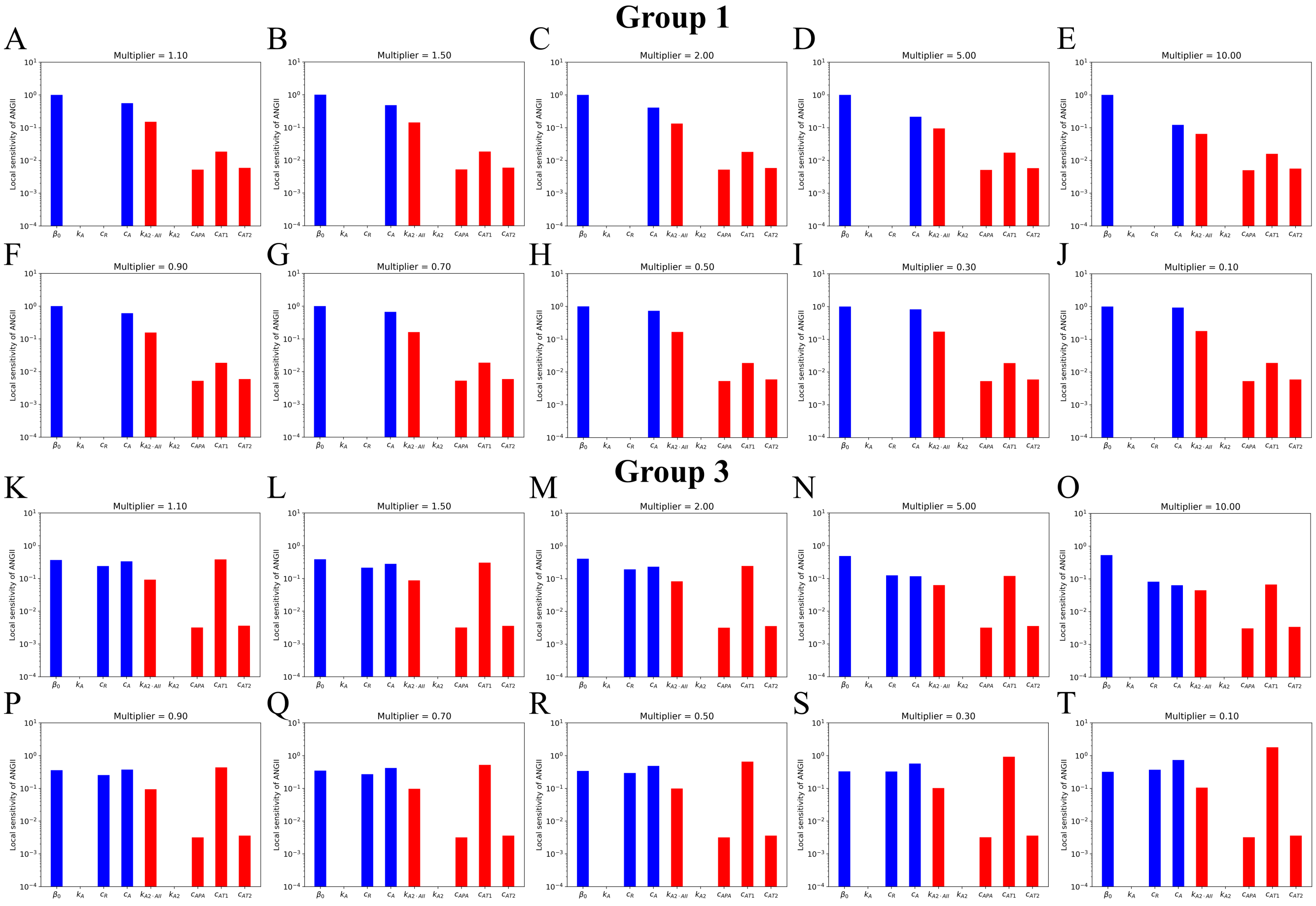


Fig. S10: Local sensitivity of the parameters for ANGII at 10 days post-infection and $ueACE2_0 = 1000$. Blue bars indicate that the output and the input change in the same direction, and the red bars indicate that the output and input change in opposite directions. (Rows 1 and 2: A–J) Group 1 hypertensive patients with no feedback from ANGII·AT1R to renin. (Rows 3 and 4: K–T) Group 3 hypertensive patients with feedback from ANGII·AT1R to renin. The parameters are multiplied by a multiplier ($m_S$). The values of $m_S$ increase across the first and third rows: (A, K) 1.1, (B, L) 1.5, (C, M) 2, (D, N) 5, (E, O) 10. Values of $m_S$ decrease across the second and fourth rows: (F, P) 0.9, (G, Q) 0.7, (H, R) 0.5, (I, S) 0.3, and (J, T) 0.1. The parameters $k_{ACE2\cdot ANGII}$ and $k_{ACE2}$ are represented as $k_{A2\cdot AII}$ and $k_{A2}$, respectively, on the $x$-axes.

### S4. Supplementary tables

Table S1: Variations of RAS peptides from homeostasis concentrations at day 10 for Group 1: hypertensive patients with no feedback from ANGII·AT1R to renin.

| $ueACE2_0$ | Sex | %Renin | %AGT | %ANGI | %ANGII | %ANG1–7 | %ACE2·ANGII | %ANGIV | %ANGII·AT1R | %ANGII·AT2R | %MAS·ANG1–7 |
|---|---|---|---|---|---|---|---|---|---|---|---|
| Older | | | | | | | | | | | |
| 200 | m | 0 | 72.86 | -100 | -100 | -100 | -100 | -100 | -100 | -100 | -100 |
| 400 | m | 0 | 72.86 | -100 | -100 | -100 | -100 | -100 | -100 | -100 | -100 |
| 600 | f | 0 | 71.91 | -95.97 | -99.92 | -99.99 | -99.99 | -99.92 | -99.92 | -99.92 | -99.99 |
| 800 | f | 0 | 63.19 | -66.44 | -93.51 | -99.14 | -99.14 | -93.51 | -93.51 | -93.51 | -99.14 |
| Middle-aged | | | | | | | | | | | |
| 1000 | m | 0 | 49.91 | -39.65 | -74.68 | -92.03 | -92.03 | -74.68 | -74.68 | -74.68 | -92.03 |
| 1200 | f | 0 | 38.60 | -25.38 | -56.52 | -79.55 | -79.55 | -56.52 | -56.52 | -56.52 | -79.55 |
| Younger | | | | | | | | | | | |
| 1400 | m | 0 | 32.48 | -19.54 | -46.77 | -70.50 | -70.50 | -46.77 | -46.77 | -46.77 | -70.50 |
| 1600 | m | 0 | 29.80 | -17.28 | -42.55 | -66.04 | -66.04 | -42.55 | -42.55 | -42.55 | -66.04 |
| 1800 | f | 0 | 29.32 | -16.90 | -41.81 | -65.23 | -65.23 | -41.81 | -41.81 | -41.81 | -65.23 |
| 2000 | f | 0 | 28.41 | -16.18 | -40.40 | -63.64 | -63.64 | -40.40 | -40.40 | -40.40 | -63.64 |

Table S2: Variations of RAS peptides from homeostasis concentrations at day 10 for Group 2: normotensive patients with no feedback from ANGII·AT1R to renin.

| $ueACE2_0$ | Sex | %Renin | %AGT | %ANGI | %ANGII | %ANG1–7 | %ACE2·ANGII | %ANGIV | %ANGII·AT1R | %ANGII·AT2R | %MAS·ANG1–7 |
|---|---|---|---|---|---|---|---|---|---|---|---|
| Older | | | | | | | | | | | |
| 200 | m | 0 | 27.21 | -100 | -100 | -100 | -99.99 | -100 | -100 | -100 | -99.99 |
| 400 | m | 0 | 27.21 | -100 | -100 | -100 | -99.99 | -100 | -100 | -100 | -99.99 |
| 600 | f | 0 | 26.86 | -97.60 | -99.94 | -99.99 | -99.99 | -99.93 | -99.93 | -99.93 | -99.99 |
| 800 | f | 0 | 23.60 | -77.13 | -94.23 | -99.23 | -99.23 | -94.23 | -94.23 | -94.23 | -99.23 |
| Middle-aged | | | | | | | | | | | |
| 1000 | m | 0 | 18.64 | -52.81 | -76.19 | -92.50 | -92.50 | -76.19 | -76.19 | -76.19 | -92.50 |
| 1200 | f | 0 | 14.42 | -36.69 | -58.03 | -80.26 | -80.26 | -58.03 | -58.03 | -58.03 | -80.26 |
| Younger | | | | | | | | | | | |
| 1400 | m | 0 | 12.13 | -29.27 | -48.11 | -71.24 | -71.24 | -48.11 | -48.11 | -48.11 | -71.24 |
| 1600 | m | 0 | 11.13 | -26.25 | -43.80 | -66.78 | -66.78 | -43.80 | -43.80 | -43.80 | -66.78 |
| 1800 | f | 0 | 10.96 | -25.73 | -43.04 | -65.97 | -65.23 | -43.04 | -43.04 | -43.04 | -65.97 |
| 2000 | f | 0 | 10.61 | -24.74 | -41.59 | -64.37 | -64.37 | -41.59 | -41.59 | -41.59 | -64.37 |

Table S3: Variations of RAS peptides from homeostasis concentrations at day 10 for Group 3: hypertensive patients with feedback from ANGII·AT1R to renin.

| $ueACE2_0$ | Sex | %Renin | %AGT | %ANGI | %ANGII | %ANG1–7 | %ACE2·ANGII | %ANGIV | %ANGII·AT1R | %ANGII·AT2R | %MAS·ANG1–7 |
|---|---|---|---|---|---|---|---|---|---|---|---|
| Older | | | | | | | | | | | |
| 200 | m | 926 | 72.83 | -100 | -100 | -100 | -100 | -100 | -100 | -100 | -100 |
| 400 | m | 926 | 72.85 | -100 | -100 | -100 | -100 | -100 | -100 | -100 | -100 |
| 600 | f | 926 | 63.14 | -58.67 | -99.19 | -99.99 | -99.99 | -99.19 | -99.19 | -99.19 | -99.99 |
| 800 | f | 191 | 44.76 | -2.26 | -81.09 | -97.50 | -99.14 | -93.51 | -93.51 | -93.51 | -99.14 |
| Middle-aged | | | | | | | | | | | |
| 1000 | m | 67 | 34.41 | 1.12 | -57.57 | -86.64 | -86.64 | -57.57 | -57.57 | -57.57 | -86.64 |
| 1200 | f | 35 | 26.30 | 1.40 | -40.92 | -72.22 | -79.55 | -56.52 | -56.52 | -56.52 | -79.55 |
| Younger | | | | | | | | | | | |
| 1400 | m | 25 | 22.03 | 1.28 | -32.99 | -62.87 | -62.87 | -32.99 | -32.99 | -32.99 | -62.87 |
| 1600 | m | 22 | 20.17 | 1.19 | -29.71 | -58.46 | -58.46 | -29.71 | -29.71 | -29.71 | -58.46 |
| 1800 | f | 21 | 19.85 | 1.18 | -29.15 | -57.66 | -57.66 | -29.15 | -29.15 | -29.15 | -57.66 |
| 2000 | f | 20 | 19.22 | 1.15 | -28.07 | -56.12 | -56.12 | -28.07 | -28.07 | -28.07 | -56.12 |

Table S4: Variations of RAS peptides from homeostasis concentrations at day 10 for Group 4: normotensive patients with feedback from ANGII·AT1R to renin.

| $ueACE2_0$ | Sex | %Renin | %AGT | %ANGI | %ANGII | %ANG1–7 | %ACE2·ANGII | %ANGIV | %ANGII·AT1R | %ANGII·AT2R | %MAS·ANG1–7 |
|---|---|---|---|---|---|---|---|---|---|---|---|
| Older | | | | | | | | | | | |
| 200 | m | 481 | 27.21 | -100 | -100 | -100 | -99.99 | -100 | -100 | -100 | -99.99 |
| 400 | m | 481 | 27.21 | -100 | -100 | -100 | -99.99 | -100 | -100 | -100 | -99.99 |
| 600 | f | 481 | 25.16 | -86.06 | -99.62 | -99.99 | -99.99 | -99.62 | -99.62 | -99.62 | -99.99 |
| 800 | f | 352 | 10.90 | 3.58 | -73.84 | -96.53 | -96.53 | -73.84 | -73.84 | -73.84 | -96.53 |
| Middle-aged | | | | | | | | | | | |
| 1000 | m | 122 | 8.19 | 4.79 | -47.13 | -83.35 | -83.35 | -47.13 | -47.13 | -47.13 | -83.35 |
| 1200 | f | 63 | 6.23 | 3.8 | -31.18 | -67.64 | -67.64 | -31.18 | -31.18 | -31.18 | -67.64 |
| Younger | | | | | | | | | | | |
| 1400 | m | 45 | 5.22 | 3.15 | -24.32 | -58.06 | -58.06 | -24.32 | -24.32 | -24.32 | -58.06 |
| 1600 | m | 39 | 4.78 | 2.86 | -21.61 | -53.67 | -53.67 | -21.61 | -21.61 | -21.61 | -53.67 |
| 1800 | f | 38 | 4.70 | 2.81 | -21.15 | -52.89 | -52.89 | -21.15 | -21.15 | -21.15 | -52.89 |
| 2000 | f | 36 | 4.56 | 2.72 | -20.28 | -51.37 | -51.37 | -20.28 | -20.28 | -20.28 | -51.37 |

Table S5: Percentage increase in unconverted ANGII at 10 days post-infection. Group 1: hypertensive patients with no feedback from ANGII·AT1R to renin. Group 2: normotensive patients with no feedback from ANGII·AT1R to renin. Group 3: hypertensive patients with feedback from ANGII·AT1R to renin. Group 4: normotensive patients with feedback from ANGII·AT1R to renin.

| $ueACE2_0$ | Sex | Group 1 | Group 2 | Group 3 | Group 4 |
|---|---|---|---|---|---|
| Older | | | | | |
| 200 | m | 100 | 100 | 100 | 100 |
| 400 | m | 100 | 100 | 100 | 100 |
| 600 | f | 99.99 | 99.99 | 99.99 | 99.99 |
| 800 | f | 99.14 | 99.23 | 97.50 | 96.54 |
| Middle-aged | | | | | |
| 1000 | m | 92.03 | 92.50 | 86.64 | 83.35 |
| 1200 | f | 79.55 | 80.26 | 72.22 | 67.64 |
| Younger | | | | | |
| 1400 | m | 70.50 | 71.24 | 62.86 | 58.06 |
| 1600 | m | 66.04 | 66.78 | 58.46 | 53.67 |
| 1800 | f | 65.23 | 65.97 | 57.66 | 52.89 |
| 2000 | f | 63.64 | 64.37 | 56.12 | 51.37 |
